# Fronto-temporal coupling dynamics during spontaneous activity and auditory processing

**DOI:** 10.1101/2019.12.23.886770

**Authors:** Francisco García-Rosales, Luciana Lopez-Jury, Eugenia Gonzalez-Palomares, Yuranny Cabral-Calderín, Julio C. Hechavarría

## Abstract

Most mammals rely on the extraction of acoustic information from the environment in order to survive. However, the mechanisms that support sound representation in auditory neural networks involving sensory and association brain areas remain underexplored. In this study, we address the functional connectivity between an auditory region in frontal cortex (the frontal auditory field, FAF) and the auditory cortex (AC) in the bat *Carollia perspicillata*. The AC is a classic sensory area central for the processing of acoustic information. On the other hand, the FAF belongs to the frontal lobe, a brain region involved in the integration of sensory inputs, modulation of cognitive states, and in the coordination of behavioural outputs. The FAF-AC network was examined in terms of oscillatory coherence (local-field potentials, LFPs), and within an information theoretical framework linking FAF and AC spiking activity. We show that in the absence of acoustic stimulation, simultaneously recorded LFPs from FAF and AC are coherent in low frequencies (1-12 Hz). This “default” coupling was strongest in deep AC layers and was unaltered by acoustic stimulation. However, presenting auditory stimuli did trigger the emergence of coherent auditory-evoked gamma-band activity (>25 Hz) between the FAF and AC. In terms of spiking, our results suggest that FAF and AC engage in distinct coding strategies for representing artificial and natural sounds. Taken together, our findings shed light onto the neuronal coding strategies and functional coupling mechanisms that enable sound representation at the network level in the mammalian brain.

## 1 Introduction

Many animals rely on the processing of acoustic information for survival. Nevertheless, the mechanisms by which sounds are represented in neural networks involving distant areas in the brain remain obscure. In the mammalian cortex, sensory and integration areas have been described as part of putative neural networks tasked with sound processing. The auditory cortex (AC), for example, plays an important role in sound analysis and even in coordinating acoustically guided behaviours (Song et al., 2010; Li et al., 2017). Neuronal activity within the AC represents a large range of acoustic properties including spectrotemporal structure (Gaese and Ostwald, 1995; Lu et al., 2001; Yin et al., 2011; Gaucher et al., 2013; Kanold et al., 2014; Gao and Wehr, 2015; Lu et al., 2016; Martin et al., 2017; Sheikh et al., 2019), sound source location encompassing azimuth/elevation coding (Recanzone, 2000; Mrsic-Flogel et al., 2005; Salminen et al., 2015; Trapeau and Schonwiesner, 2018) and target distance processing (Suga and O’Neill, 1979; Hechavarria et al., 2013; Bartenstein et al., 2014; Beetz et al., 2016), as well as abstract properties such as sound “emotional valence” (Concina et al., 2019) and future behavioural outcomes based on auditory stimuli (Francis et al., 2018).

Regions within the frontal lobe of the mammalian brain also participate in auditory processing and could in principle synchronize their activity with that of canonical auditory areas, such as the AC (see (Winkowski et al., 2018)). Frontal and AC regions are strongly connected through feedforward and feedback anatomical pathways (Kobler et al., 1987; Medalla and Barbas, 2014; Plakke and Romanski, 2014; Winkowski et al., 2018). Neurons within the prefrontal cortex (PFC, a region in the frontal lobe) respond to sounds when the latter possess rich spectrotemporal dynamics (Eiermann and Esser, 2000; Kanwal et al., 2000; Romanski and Goldman-Rakic, 2002). Additionally, PFC is thought to engage in cognitive processes ranging from attention, learning, and memory formation/retrieval, to decision making (Miller, 2000; Floresco and Ghods-Sharifi, 2007; St Onge et al., 2011; Gourley et al., 2013; Gilmartin et al., 2014; Pezze et al., 2014; Helfrich and Knight, 2016; Kim et al., 2016; Werchan et al., 2016; Helfrich et al., 2017). This area could thus be a fundamental node for sound evaluation in auditory networks, and even for the implementation of acoustically guided behaviours.

Though there is increasing evidence supporting the idea of frontal-AC functional networks for auditory processing, specifics regarding activity coupling within this network remain unknown. For example, it remains unclear if different types of neural signals (i.e. spikes and local field potentials, LFPs) measured simultaneously in frontal and AC areas synchronize during spontaneous activity and during listening. Moreover, it is unknown whether frontal activity displays preferential coupling patterns with certain layers of the AC. Assessing the latter can only be achieved by conducting simultaneous measurements from frontal and AC regions using layer-specific intracranial recordings to study spikes and LFPs.

In the current study, we address the functional connectivity in a fronto-auditory cortical circuit in bats (species *Carollia perspicillata*). The bat AC has been studied extensively, and it has been shown that oscillatory and spiking activity patterns in the bat cortex are in accordance with those observed during the processing of artificial and naturalistic sounds in other animal models, including speech in humans. This comprises phenomena such as multiscale temporal processing of acoustic streams (Giraud and Poeppel, 2012; Hyafil et al., 2015; Hechavarria et al., 2016b; Teng et al., 2017; Garcia-Rosales et al., 2018a), interactions between spikes and LFPs for audition (Lakatos et al., 2005; Kayser et al., 2009; Arnal and Giraud, 2012; Kayser et al., 2012; Gilmartin et al., 2014; Garcia-Rosales et al., 2018b; Garcia-Rosales et al., 2019), and gamma-band activity for communication call processing (Medvedev and Kanwal, 2008). In bats, there exists a region within the frontal lobe that is responsive to sounds: the frontal auditory field (FAF; (Kobler et al., 1987; Eiermann and Esser, 2000; Kanwal et al., 2000)). This region is anatomically connected with the AC (Kobler et al., 1987), but it also receives auditory afferents via a non-lemniscal pathway through the suprageniculate nucleus of the thalamus, bypassing main auditory centres in the midbrain (Kobler et al., 1987). In addition to pure tones, neurons in the FAF encode spectrotemporally complex sounds with variable latencies and response properties (Eiermann and Esser, 2000; Kanwal et al., 2000; Lopez-Jury et al., 2019). Our goal was to examine specifics of fronto-AC activity in awake bats during the processing of acoustic streams. We tackled this question by quantifying neural synchronization in the FAF-AC network in terms of oscillatory coherence (a mechanism underlying interareal communication; (Fries, 2015)), during both spontaneous activity and the processing of natural and artificial acoustic sequences.

We found that the FAF-AC network is synchronized by default (i.e. without sensory stimulation) in low-frequencies (up to 12 Hz), and that coherence with the FAF is strongest in deep laminae of the AC. In addition, low-frequency coherence between the two structures remains unchanged during acoustic processing, and auditory-evoked gamma-band synchronization emerges at stimulus onset without clear layer specificity. Finally, based on an information theoretical framework, our data suggest that the neuronal coding of acoustic streams in FAF and AC may occur with non-overlapping neural codes. Taken together, the data presented in this manuscript offer insights into the strategies for sound representation in fronto-AC neural networks.

## 2 Results

### 2.1 Stimulus-related LFPs in AC lag relative to those in FAF

We recorded electrophysiological data from the primary AC, paired with penetrations from the FAF, in 5 awake *Carollia perspicillata* bats (all males; n = 50 penetrations). Recordings in the AC were performed with laminar electrodes inserted perpendicularly into the brain, spanning depths of 0-750μm as measured from the cortical surface. Each penetration in AC was paired with a simultaneous recording from the FAF using a single carbon electrode at an average depth of 313 ± 56 μm (mean ± std). Auditory stimuli consisted of artificially constructed syllabic trains with repetition rates of 5.28 and 97 Hz in order to test for slow and fast periodicities in acoustic streams, plus another syllabic train that had no clear rhythmicity as syllables were presented in a Poisson-like sequence with 70 Hz average rate (see Methods). The trains consisted of a repeated short duration, broadband distress syllable from *C. perspicillata*, recorded in previous work (Hechavarria et al., 2016a), whose spectrotemporal design is typical of distress syllables emitted by this species. In addition, we presented a natural distress vocalization (*“nat*” throughout the text) that has been used in previous research (Hechavarria et al., 2016b; Garcia-Rosales et al., 2018a; Garcia-Rosales et al., 2019), and which comprises temporal modulations in low (ca. 4 Hz) and high-frequency ranges (> 50 Hz), corresponding to the bout and syllabic periodicities, respectively, typical of this animal’s distress vocalizations (Hechavarria et al., 2016a).

Grand average traces of simultaneously recorded LFPs from FAF and AC (the latter at a depth of 450μm, corresponding to input layers in AC) during acoustic stimulation are shown in **Fig. 1A**. LFP responses from frontal and auditory cortical regions showed clear modulation by the acoustic streams that were well-correlated across structures for each stimulus tested, particularly at depths > 200 μm (Fig. S1). Remarkably, population averaged LFPs appeared “faster” in the FAF than in the AC (see **Fig 1A**, right column; blue: FAF; orange: AC at 450 μm) relative to stimulus onset. In fact, cross-correlation analyses of the traces depicted in **Fig. 1A** showed that LFPs recorded in frontal regions preceded those recorded from the AC by at least 4 ms (**Fig. 1B**), effectively indicating that primary auditory cortical stimulus-related LFPs in input layers “lag” relative to those in the FAF. We confirmed this trend by systematically determining the temporal lag between LFPs in the AC at various depths, and LFPs from the FAF. **Figure 1C** summarizes the results by illustrating, for each stimulus, the distribution of lags between FAF and AC across depths (note that negative lags indicate FAF “leading”). Indeed, we observed a significant effect of LFPs in the FAF being “faster” than those in the AC, robust across depths in the latter structure, and also across stimuli (**Fig 1C**; FDR-corrected tailed Wilcoxon signed rank tests, testing that the medians of the lag distributions were significantly less than 0 across electrodes; p_corr_ < 0.05; precise p_corr_ values are indicated next to each heatmap). These results were corroborated by testing FAF-AC lags in LFPs but considering only LFP pairs that were well correlated across structures (correlation coefficients higher than 0.5, shown in Fig. S1).

**Fig. 1.**
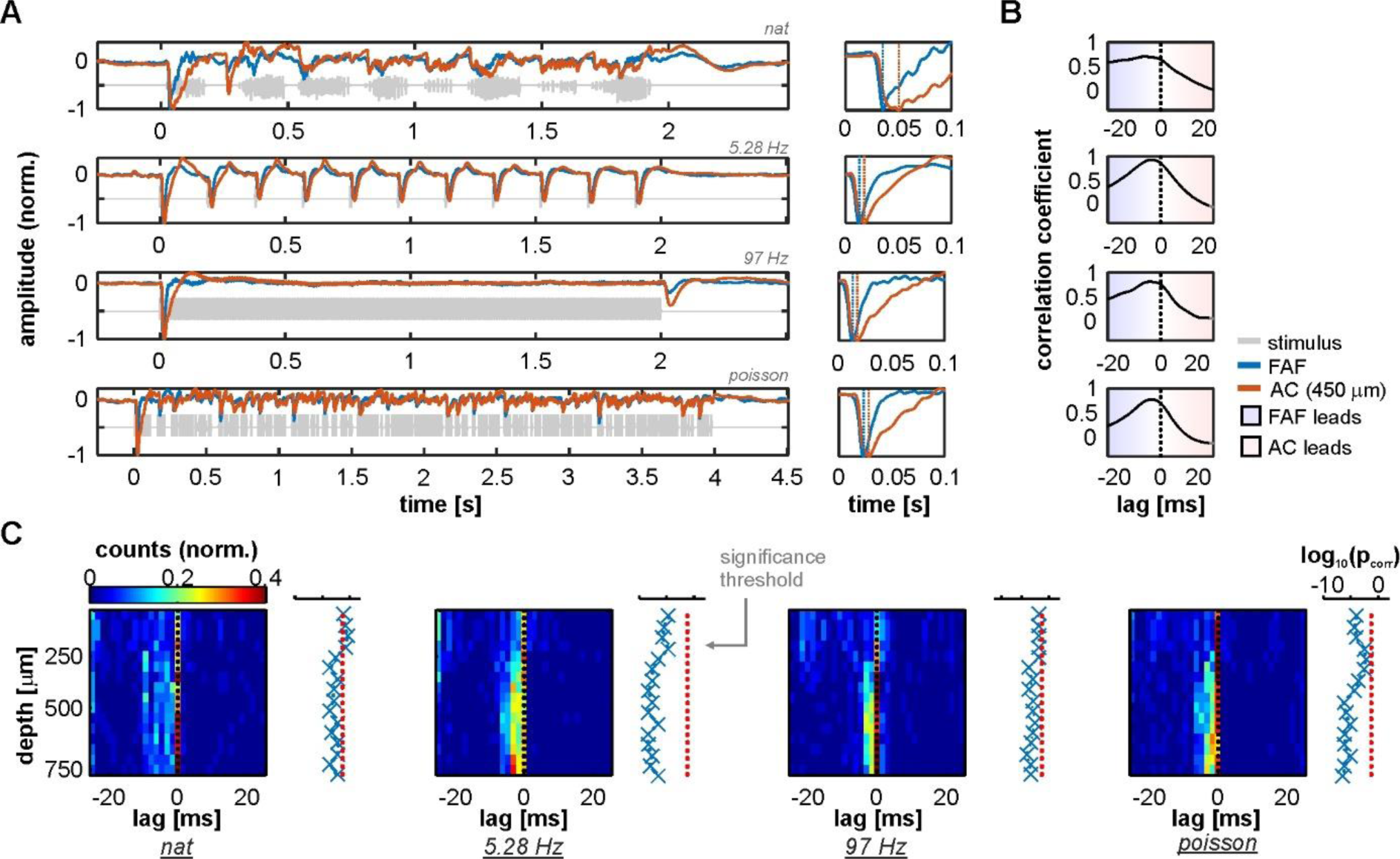
LFP stimulus-related activity in the frontal auditory field (FAF) precedes that of the auditory cortex (AC). (**A**) *Left column:* Grand average across all 50 penetrations of LFPs recorded from the FAF (blue) and the AC at a depth of 450 μm (orange) in response to the four stimuli tested (ordered from top to bottom: natural sequence (*nat*), 5.28 Hz train, 97 Hz train, and the Poisson syllabic sequence (*poisson*); grey traces). *Right column*: zoom into the first 100 ms after stimulus onset. Negative peaks in the evoked potential are marked with vertical dashed lines. Note that peaks in the FAF occur earlier than in the AC. (**B**) Cross-correlation between traces in A (*left column*). Peaks in negative lags indicate that FAF field-potentials lead those in the AC. (**C**) Peak lags between the cross-correlation of FAF LFPs and AC LFPs, for all penetrations (n = 50) and across recording depths. Next to each heatmap, log-scaled corrected p values testing that the peak distribution is significantly below 0 (FDR corrected tailed Wilcoxon signed rank tests; p_corr_ < 0.05 for significance).

That LFPs in the frontal auditory field “lead” relative to those in the auditory cortex suggests the presence of fast inputs into frontal auditory areas, agreeing with a non-lemniscal auditory pathway converging into the FAF and consisting of as few as four synapses in bats (Kobler et al., 1987). We sought for evidence of fast neuronal responses in FAF that would support these observations by measuring response latencies of the neuronal spiking recorded simultaneously in both structures (see Methods). We observed that spiking responses from the FAF could in fact be as fast as spiking responses from the AC, although this effect was found only in a subpopulation of neurons (on average 5.65% ± 3.19% of the units considered, across stimuli and channels; see **Fig. 2A, B**, which depicts example responses from one FAF and one AC unit at 450 μm recorded simultaneously).

**Fig. 2.**
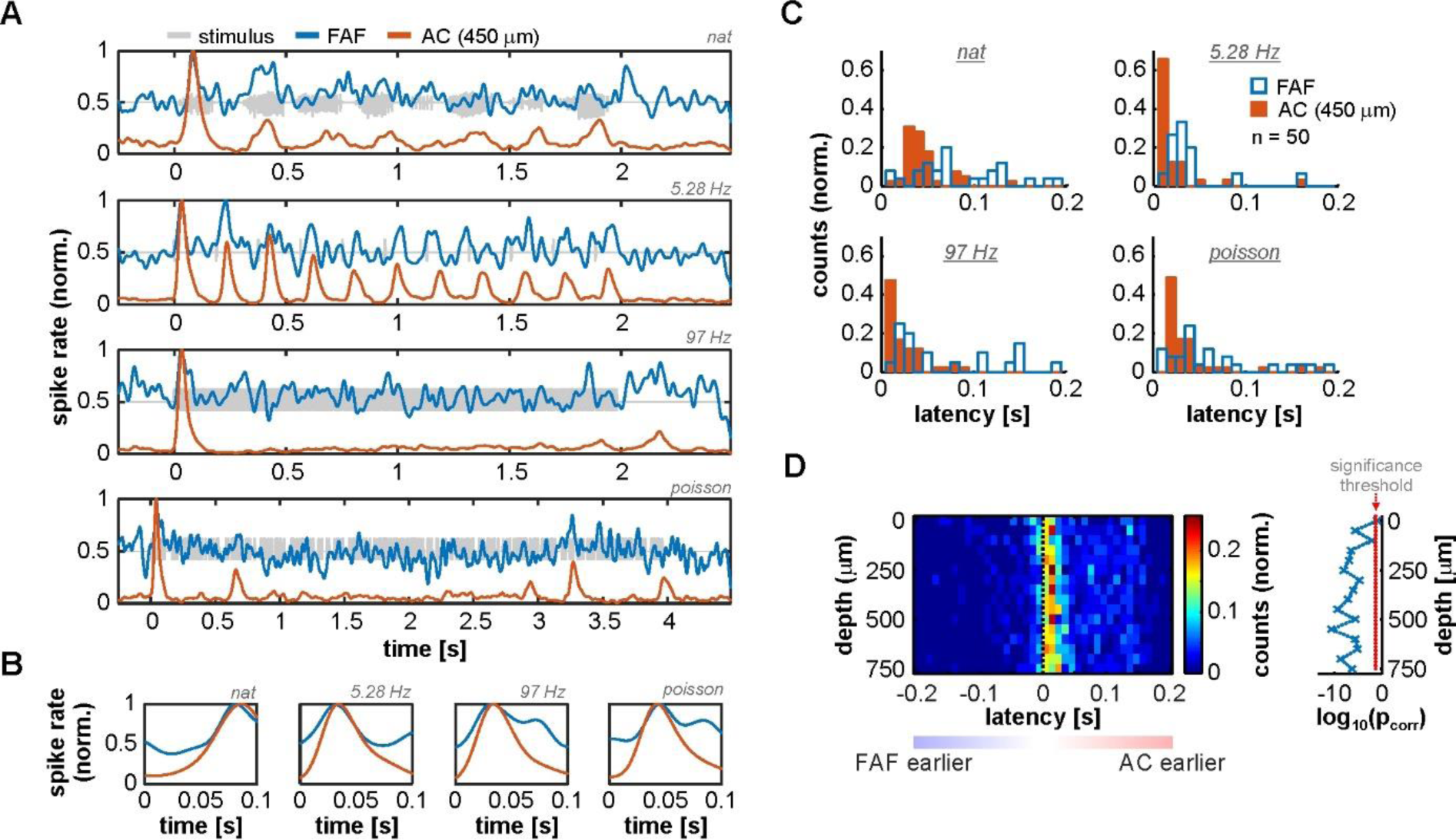
Spiking activity suggests the presence of fast inputs into the FAF. (**A**) Spiking responses from two simultaneously recorded units in the FAF (blue) and the AC (at 450 μm; blue), in response to all stimuli tested (top to bottom). (**B**) Zoom-in into the first 100 ms after stimulus onset of the examples shown in **A**. Note that, for this pair, the peak response for the FAF unit was at least as fast as the for auditory cortical one. (**C**) Latency distribution of FAF (blue) units and AC units at 450 μm, for all stimuli (n = 50 penetrations). The FAF was, overall, sluggish in comparison to the AC. (**D**) Response latency difference between simultaneously recorded spiking for FAF and AC at different depths (positive difference, FAF slower than AC; negative difference indicates the opposite). In some cases, FAF spiking responses occurred earlier than AC responses, although the AC was in general significantly faster than the FAF across channels, except in the case of the most superficial contact (FDR-corrected Wilcoxon signed rank tests, significance when p_corr_ < 0.05). Log-converted p values and significance threshold are shown to the right of the latency distributions. The threshold is indicated as a red dashed line.

Measured neuronal response latencies from both structures yielded that AC spiking was on average faster than the FAF spiking (**Fig. 2C**; for illustrative purposes AC responses are those recorded at 450 μm). Still, some FAF units exhibited response latencies below 10 ms, indicative of fast acoustic inputs into this structure. By subtracting latencies from simultaneously recorded units in the FAF and the AC (across depths; latencies were pooled from all tested stimuli, but paring was only done within a particular stimulus), it became evident that auditory cortical spiking was typically faster than its FAF counterpart, although some latencies from frontal regions were shorter than those in the AC. **Figure 2D** shows the distribution of latency differences (across cortical depths in the AC), depicting the abovementioned observations. Latency differences were significantly higher than 0 for all recording depths in the AC, except in the case of the most superficial channel (FDR-corrected Wilcoxon signed rank tests, significance when p_corr_ < 0.05; corrected p values across channels are given to the right of the latency distribution heatmap in **Fig. 2D**).

Altogether, these results provide evidence supporting that the FAF receives fast auditory inputs. Notwithstanding, such inputs do not necessarily elicit equally fast spiking, suggesting that the neuronal dynamics in frontal areas are “sluggish” in comparison to primary AC. Sluggish dynamics can arise from multiple factors, and may be essential for sensory integration in the frontal cortex (see Discussion).

### 2.2 FAF and AC synchronize in low frequency LFP bands during spontaneous activity

Local-field potentials recorded in the FAF and the AC typically showed visible phase synchronization, even in the absence of acoustic stimulation (see **Fig. 3A**, where LFP traces from both structures during a single 3 s epoch of spontaneous activity are depicted). We quantified phase coherence between the two structures by means of the imaginary part of the coherency (“iCoh” in this manuscript; see Methods and (Nolte et al., 2004)), for data recorded both during spontaneous and sound-driven activities. The iCoh metric allows to minimize spurious phase-synchrony attributable, for example, to common referencing and passive spreading of field potentials, by effectively removing non-lagged phase correlations (Bastos and Schoffelen, 2015).

**Fig. 3.**
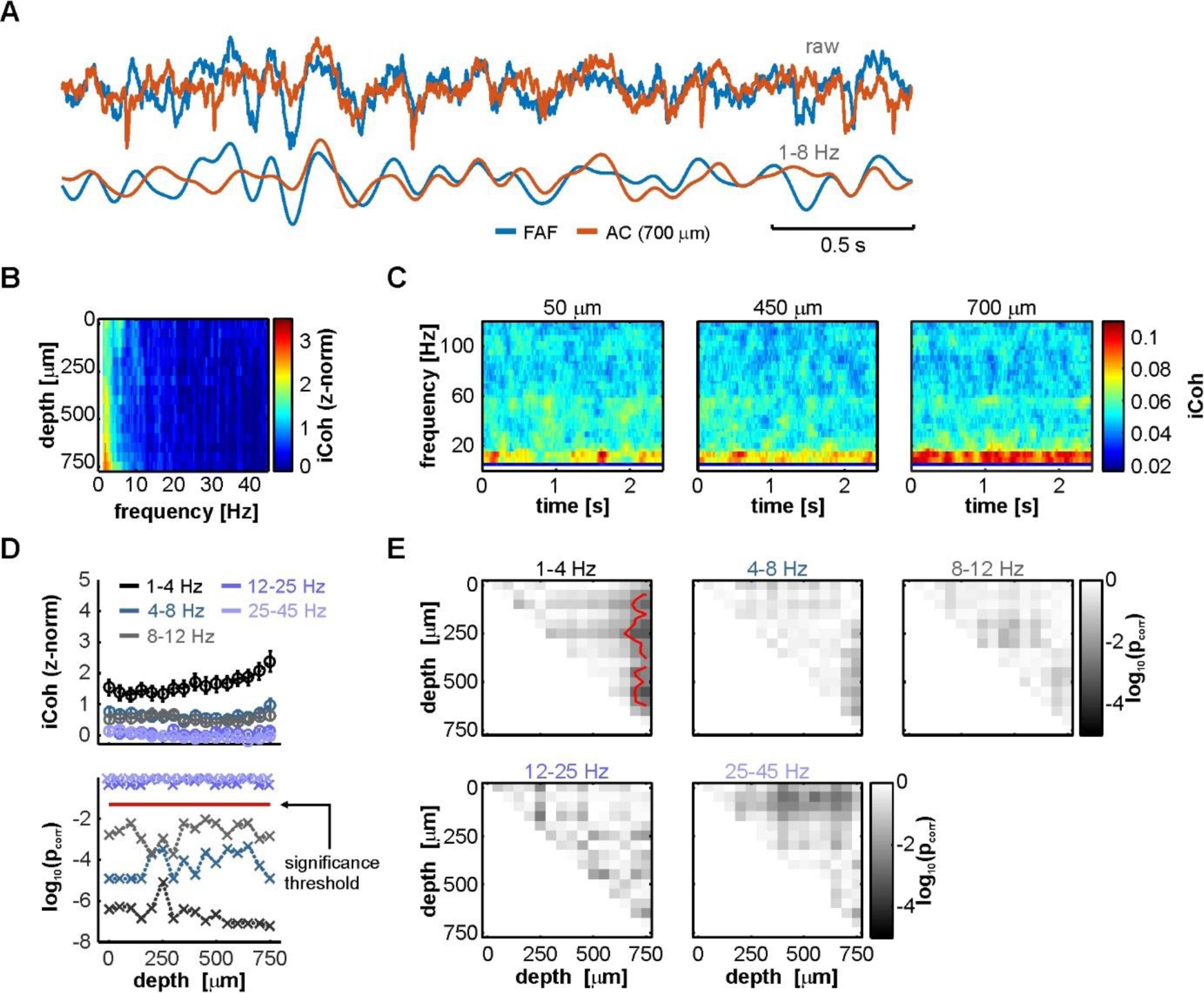
LFPs from FAF and AC are low-frequency coherent during spontaneous activity in a depth-dependent manner. (**A**) Simultaneously recorded LFP trace from the FAF (blue) and the AC at a depth of 700 μm (orange). Raw (top pair) and low-frequency filtered (bottom pair) traces are shown. (**B**) Frequency-dependent average imaginary coherence (iCoh), z-normalized to a surrogate distribution, across recording depths in the AC. Deep channels showed, on average, the strongest coherence values at low frequencies. (**C**) Time-resolved iCoh using the same segments as in **B**, with a sliding window of 200 ms (see Methods), the same used for analyzing stimulus-related synchronization. (**D**) *Top*: depth-dependent population z-normalized iCoh from **B,** for distinct frequency bands (delta, theta, alpha, beta and low gamma, see Methods; frequency ranges indicated in the plot), across all penetrations (shown as mean ± SEM). *Bottom*: log-scaled corrected p values after testing, per frequency band, whether z-normalized iCoh values were significantly higher than 0 across penetrations, per AC depth (FDR-corrected Wilcoxon signed rank tests, p_corr_ < 0.05 for significance; threshold indicated as a horizontal red dashed line). (**E**) Significance matrices comparing, per frequency band, z-normalized iCoh values across different depths. Each cell (*i, j*) in a matrix depicts the log-scaled p_corr_ obtained from statistically comparing coherence at channels with depths *i* and *j* in the AC. Red contour lines delimit regions of statistical significance (FDR-corrected Wilcoxon signed rank tests, significance when p_corr_ < 0.05).

Coherence analyses revealed that, as shown in **Fig. 3A**, FAF and AC were synchronized in low frequencies. **Figure 3B** depicts population averaged z-normalized (to a surrogate distribution where phase relationships were abolished; see Methods) iCoh values (z-iCoh) across electrode depths in the AC. Elevated low-frequency coherence is evident in deep layers of the AC, suggesting as well that FAF-AC synchrony was depth-dependent. A time-resolved analysis of iCoh (**Fig. 3C**) over the same LFP traces used to calculate values in **Fig. 3B** also showed that low-frequency phase synchrony was strongest in deeper channels, and furthermore limited to the low-frequency region of the coherence spectrum. The data depicted in **Fig. 3C** are shown here for illustrative purposes and serve as a comparison with the time-resolved coherence estimations performed on LFPs recorded during acoustic processing (see below).

To statistically corroborate our observations, we divided the coherence spectrum into canonical frequency bands encompassing delta (1-4 Hz), theta (4-8 Hz), alpha (8-12 Hz), beta (12-25 Hz) and low-gamma (25-45 Hz). We then tested if z-normalized iCoh values in each band were significantly different than 0 across the population. Because of the nature of the surrogate analyses (see Methods), non-consistent phase synchronization in the data would yield a distribution of z-iCoh statistically indistinguishable from 0. In order words, z-iCoh values significantly higher than 0 suggest consistent, population-wise phase-locking between LFPs in FAF and AC during spontaneous activity. **Figure 3D** (top) depicts the z-iCoh calculated between FAF and AC oscillations for each frequency band at various AC depths. For frequency bands between 1-12 Hz (i.e. delta to alpha), we observed significantly higher than 0 z-iCoh estimates (FDR-corrected Wilcoxon signed rank tests, p_corr_ < 0.05 for significance; log-converted p values are shown in **Fig. 3D**, bottom), which became gradually lower towards higher frequencies (note also the decay in **Fig. 3B**). Beta or gamma z-iCoh distributions were not significantly different than 0 at any cortical depth.

It was also apparent that coherence values were depth dependent in the AC, particularly in the delta band (where z-iCoh was strongest, see **Fig. 3B, D**). We tested the depth dependence of coherence by comparing the distributions of z-iCoh for all pairs of channels, across all penetrations and frequency bands. Results are summarized in significance matrices depicted in **Fig. 3E**. Each cell (*i, j*) in a matrix represents the log-converted, corrected p value (FDR Wilcoxon signed rank tests), obtained after statistically comparing FAF-AC coherence using an AC channel at depth *i,* and another at depth. *j*. In the matrices, cells within red contour lines correspond to statistically significant p_corr_ values (p_corr_ < 0.05). As shown in **Fig. 3E**, deep electrodes in the AC were significantly better synchronized with the FAF, an effect only visible in the delta band.

### 2.3 FAF-AC synchronization during acoustic processing

To quantify synchronization during acoustic processing in the FAF-AC circuit, we calculated time-resolved iCoh values in response to four acoustic stimuli (a natural call, syllabic trains of 5.28 and 97 Hz, and a syllabic train with Poisson temporal structure; see above). Population average coherograms (i.e. time-frequency representations of iCoh values) are shown in **Fig. 4A-D** for all stimuli tested, and at representative depths in the AC (50, 450 and 700 μm). The most conspicuous pattern across stimuli was the appearance of low-gamma coherence (typically in the range of 25-45 Hz), which was associated with stimulus onset (at time 0) and apparently independent of auditory cortical depth.

**Fig. 4.**
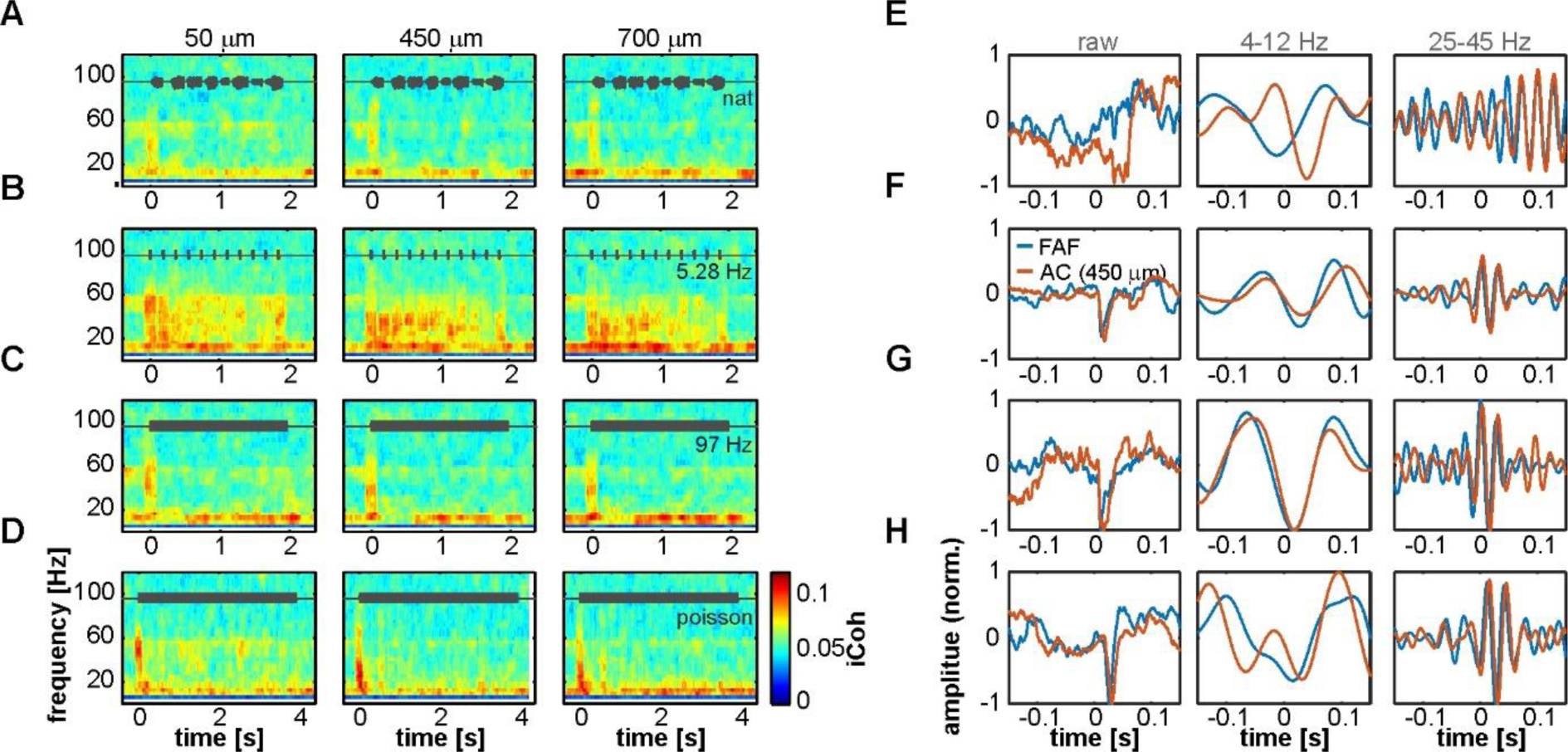
Interareal phase synchrony during acoustic sequence processing. (**A-D**) Mean time-resolved coherence between LFPs from the FAF and the AC at three representative depths (50, 450 and 700 μm), in response to the natural sequence (**A**), a syllabic train of 5.28 Hz (**B**), a syllabic train of 97 Hz (**C**), and the syllabic train with a Poisson structure (**D**). Note that low frequency synchrony is high even without acoustic stimulation, and the appearance of gamma-band evoked synchronization at the stimulus onset (time 0), albeit more weakly in response to the natural call in **A**. (**E**-**H**) LFP recordings from the AC (orange) and FAF (blue) around the time of stimulus onset (at 0 s; order in **E**-**H** corresponds to order in **A**-**D**), from single trials in a representative penetration. Left column depicts the raw LFP, whereas middle and right columns depict field-potentials filtered in 4-12 and 25-45 Hz low-frequency and gamma-bands, respectively.

Gamma synchrony was auditory-evoked, a notion strengthened when considering responses to the 5.28 Hz syllabic train (**Fig. 4B**), where the evoked coherence tracked individual syllable presentations. Remarkably, we observed little evidence for an increase of low-frequency synchrony when compared to spontaneous activity (compare heatmaps in **Fig. 4** with **Fig. 3C**). That is, even though low-frequency coherence was present before sound presentation (in line with our results using spontaneous LFPs), it did not change visibly after stimulus onset. To illustrate the occurrence of low and high frequency coherence, **Figure 4E-H** depict single-trial LFPs from a representative pair of simultaneous penetrations in the AC (450 μm depth) and FAF. Raw LFPs are shown in their broadband form (0.1-300 Hz), and filtered in frequency ranges of 4-12 Hz and 25-45 Hz (i.e. low and high frequencies oscillations). The range 4-12 Hz was chosen for low-frequency activity because spectral parameters at lower frequencies could not be reliably estimated with the window size chosen for time-resolved coherence analyses (200 ms; see Methods). Note that the occurrence of synchronized waves in the AC and FAF is clear after stimulus onset (0-ms mark).

The defined low- and high-frequency ranges were then used to quantify changes from spontaneous to stimulus-driven coherence in the FAF-AC network. A systematic, time-resolved analysis of low-frequency synchrony revealed that, across stimuli and auditory cortical depths, there was little change (calculated as percentage increase from spontaneous to sound-driven activity: [iCoh_stim_ – iCoh_spont_]/ iCoh_spont_* 100) in coherence between both structures (**Fig. 5A, D, G, J**, top heatmaps). The data showed that low frequency synchrony preceded stimulus presentation in deep layers, seldom reached 50% increase from spontaneous across stimuli (black contour lines in **Fig. 5A, D, G, J**), and when it did, it typically happened at middle AC depths. We did not observe statistical evidence showing significant increase of low frequency coherence at stimulus onset (i.e. first 100 ms after sound presentation; **Fig. 5B, E, H, K**, black traces; FDR-corrected Wilcoxon signed rank tests, significance when p_corr_ < 0.05).

**Fig. 5.**
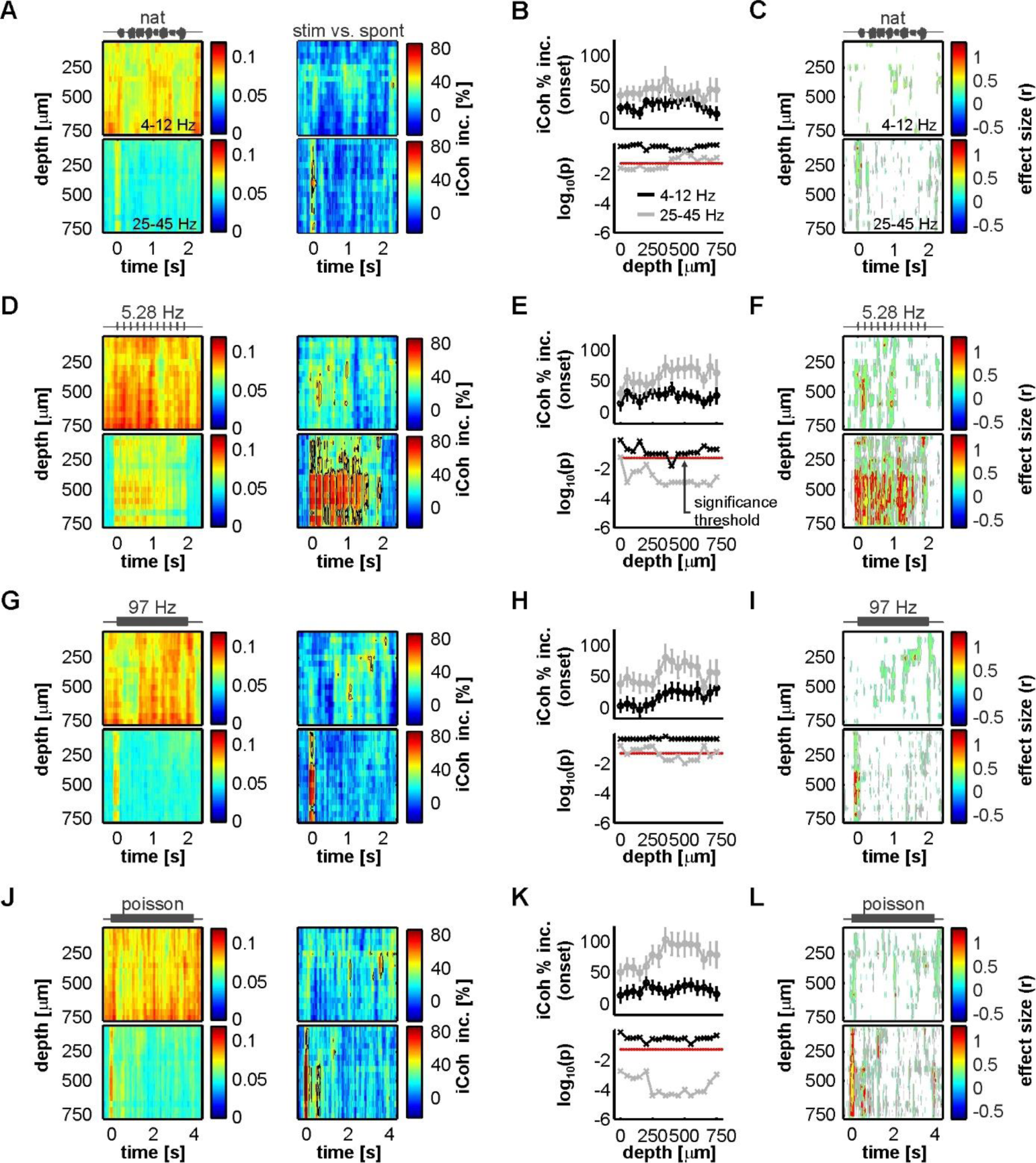
Acoustic stimulation alters FAF-AC coherence mostly in the gamma-band. (**A**) *Top*: average time course of iCoh while animals listened to the natural sequence (left) across recording depths for low frequencies (4-12 Hz), and percentage increase of coherence in that range relative to the spontaneous activity (right). *Bottom*: same as *Top*, but for iCoh values in the gamma range (25-45 Hz). Black contour lines delimit regions with average increase of coherence > 50%. (**B**) Population onset-related iCoh increase (median in the period of 0-150 ms after stimulus onset) across depths in the AC (black traces, low-frequency band iCoh; grey traces, gamma-band iCoh; shown as mean +-SEM). On the bottom subpanel, log-scaled corrected p values obtained after testing that such increase was significantly different from 0% (black, low-frequency band; grey, gamma-band; FDR-corrected Wilcoxon signed rank tests, significance when p_corr_ < 0.05, indicated as a red dashed line). (**C**) Time-resolved effect size of population iCoh percentage increase (r; see Methods) for the low-frequency band (top) and the gamma-range (bottom). Grey contour lines delimit regions of r > 0.3, whereas red contours mark regions of r > 0.5 (medium and large effect sizes, respectively). r values are only shown for time points, across channels, where the coherence increase was significantly higher than 0% (Wilcoxon signed rank test, p < 0.05). (**D-F**) Same as **A-C** but considering a syllabic train at 5.28 Hz as stimulus. (**G-I**) Same as **A-C**, the stimulus being a syllabic train at 97 Hz. (**J-L**) Same as **A**-**C**, except the stimulus was the Poisson syllabic train.

**Figure 5C, F, I, L** (top heatmaps) illustrate effect size calculations (r; see Methods) for the low-frequency coherence increase in a time resolved manner across channels. In the heatmaps, only time points where the increase was significantly different from 0 (uncorrected Wilcoxon singed-rank test, p < 0.05) are shown. From this analysis the following was evident: (i) the pattern of significance was inconsistent across stimuli for low frequency coherence; and (ii) effect sizes were typically small, with areas of medium effect size (those within grey contour lines) appearing also with an inconsistent pattern across sounds. Large effect sizes (within red contour lines) were overall only observed in small clusters, lacking consistency throughout the stimulus set. The boundaries between small, medium and large effect sizes were defined as follows: r < 0.3, small; 0.3 ≤ r < 0.5, medium; r ≥ 0.5, large (Fritz et al., 2012). Altogether, these results corroborate a lack of reliable increase in low-frequency coherence between FAF and AC during passive listening, compared to spontaneous activity.

High frequency FAF-AC coherence was considerably more sensitive to acoustic stimulation. As expected from the data depicted in **Fig. 4**, we observed a strong increase of low-gamma interareal synchronization associated to the stimulus onset (**Fig. 5A, D, G, J**, bottom heatmaps). Stimulus-evoked gamma synchrony was typically higher than 50% (reaching values as high as 80%) and tracked the syllable presentations of the 5.28 Hz syllabic train (**Fig. 5D**, bottom). Indeed, the increase of low-gamma synchrony at stimulus onset was significantly above 0 for all stimuli (**Fig. 5B, E, H, K**, grey traces; same statistical analysis as for low-frequencies; p_corr_ < 0.05), and more reliably so for channels located in input layers of the AC (in this context, electrodes at depths of 250-350 μm; layers III-IV in *C. perspicillata*’s AC span depths of 200-450 μm, see (Garcia-Rosales et al., 2019)). For the 5.28 Hz and the Poisson syllabic trains, the onset-related increase in low-gamma coherence occurred essentially along all AC depths studied. In terms of effect size (**Fig. 5C, F, I, L**; bottom heatmaps), we observed large effects of increased low-gamma coherence at stimulation onset across stimuli (less clearly in the case of the natural vocalization), with sustained, seemingly periodic increases along the time-course of the 5.28 Hz syllabic sequence. Taken together, coherence analysis results indicate that acoustic stimulation elicits auditory-evoked low-gamma synchronization between the frontal auditory field and the auditory cortex.

### 2.4 Gamma-band activity in FAF and AC

Previous studies showed the occurrence of gamma-band activity in the AC of primates, bats, and rats (Brosch et al., 2002; Medvedev and Kanwal, 2008; Vianney-Rodrigues et al., 2011). These studies reported auditory cortical gamma which was not time-locked to the onset of a stimulus, and that appeared even hundreds of milliseconds after sound presentation. Given the nature of the coherence analyses performed here, it is possible that the presence of non-locked gamma oscillations could have been overlooked in the FAF and the AC of *C. perspicillata*. To explore the occurrence of these rhythms in our dataset, we focused on the onset period of the 5.28 Hz syllable train as it was the stimulus that permitted the analysis of an onset window without the influence of subsequent sounds (after the first syllable presentation), for a sufficiently long time-lapse of at least 180 ms. The time period around the first syllable presentation in the 5.28 Hz train was subdivided into three segments: (1) a window of 90 ms spanning times before stimulus onset (*pre*); (2) a window of 90 ms starting at stimulus onset (*onset*); (3) a window of 90 ms starting 90 ms past stimulus onset (*late*); and (4) a window of 180 ms starting at stimulus onset (*full*). The span of these segments is illustrated in **Fig. 6A** together with representative LFP traces from a penetration pair. The segments were chosen in order to contrast LFP power at different frequency bands (low frequencies, 0-15 Hz; low-gamma, 25-45 Hz; high-gamma 45-80 Hz; and broad gamma 25-80 Hz) with spontaneous LFP power before stimulus presentation (*pre* window).

**Fig. 6.**
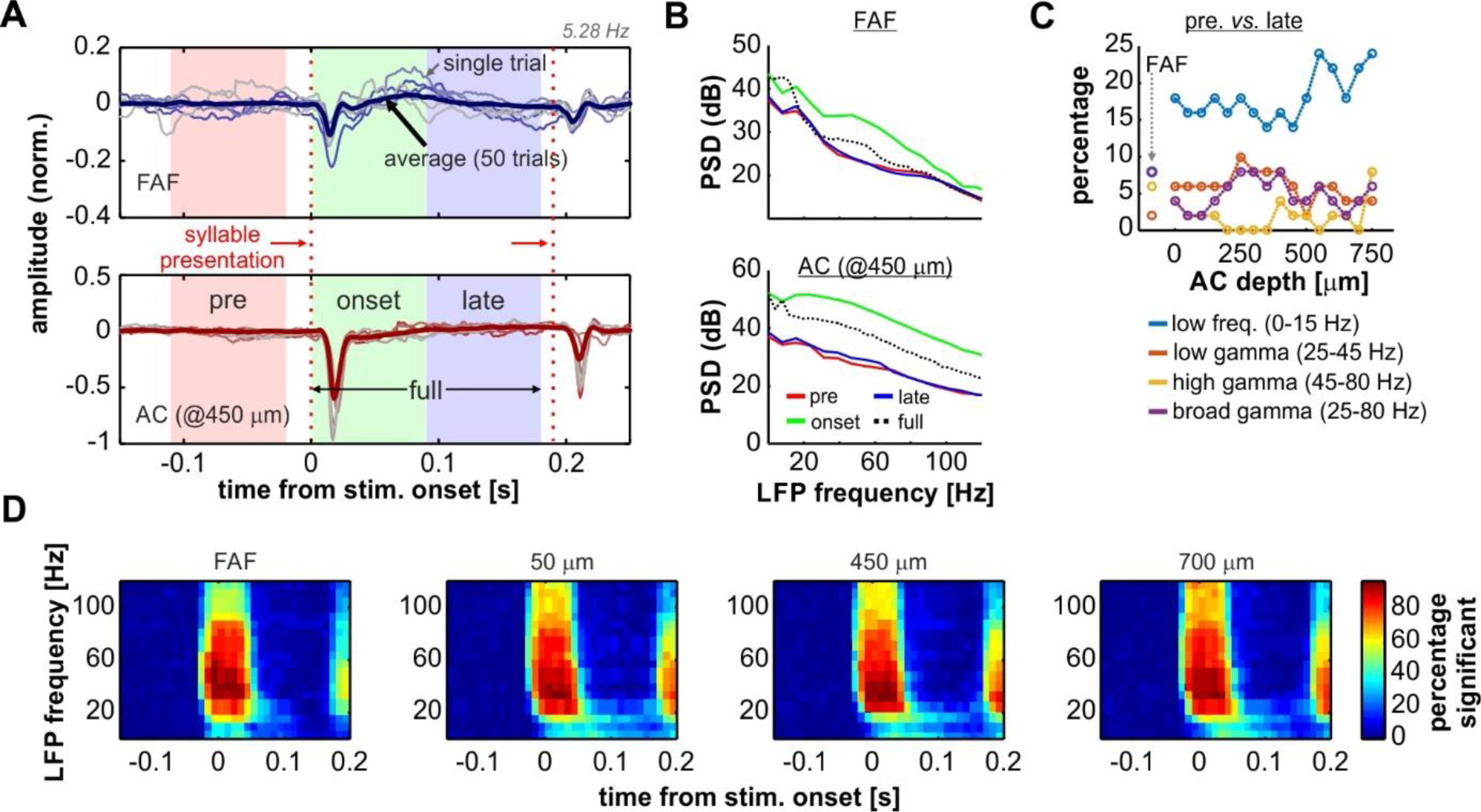
Onset related power increase in AC and FAF. (**A**) Representative LFP recordings of one penetration pair (FAF, top; AC at 450 μm, bottom), depicting 6 single trials for illustrative purposes (thin lines) and the average across all 50 trials. The time segments of *pre*, *onset*, *late*, and *full*, used for analyses (see main text), are indicated in the graphs. First and second syllable presentations of the 5.28 Hz train are indicated with vertical, red dashed lines. (**B**) Power spectral density (PSD) of the *pre* (red), *onset* (green), *late* (blue) and *full* (black, dashed) periods from the data depicted in **A** (average over the 50 trials), in the FAF (top) and AC (bottom). (**C**) Percentage of penetrations (after a total of 50) for which the power (at several frequency bands, indicated in the figure) was significantly different during the *late* period than during the *pre* period. (**D**) Time-frequency analysis illustrating the percentage of penetrations in FAF and AC (at three representative depths: 50, 450 and 700 μm) in which the power at a given time window was significantly higher than the power at a window preceding the stimulus onset.

There was a consistent increase of onset-related gamma activity in FAF and AC, particularly during the *onset* period, which was potentially linked to an evoked activation in cortex as it was associated with a broadband increase in LFP power (see **Fig. 6B**, green traces, and **Fig. S2A**). To uncover the presence of gamma within later time periods in our data, the power of different frequency bands in the *late* period was statistically compared (Wilcoxon signed rank test, significance threshold at p = 0.01; see Methods) with the power in the *pre* period on a trial-by-trial basis, per penetration (**Fig. 6C**). The percentage of penetrations in FAF where there was a power increase in low frequencies was of 8%, reaching between 14-24% in the AC. We also observed a relatively small number of penetrations (<10%) either in AC or FAF in which there was a significant power increase in gamma for the *late* period as compared to the *pre* segment. We further calculated the time course of the percentage of penetrations showing significant power increase at times surrounding the stimulus presentation for both FAF and AC, at various LFP frequencies (**Fig. 6D**; see Methods). As expected from the data shown in **Fig. 6B-C** and **Fig. S2A**, up to about 50 ms after stimulus onset there was a high percentage of penetrations (ca. 80%; cf. with **Fig. S2A**) where the power in gamma increased significantly in either structure. This number was relatively low (< 20%) for times beyond 50-60 ms after stimulus onset.

The FAF-AC circuit exhibited increased auditory-evoked gamma band coherence, related to the onset of acoustic stimulation (**Figs. 4, 5**). We tested to what extent we could disentangle gamma activity in our data from a non-specific broadband response, by means of previously used approach which relies on comparing the relative power distributions of gamma and low frequency LFPs (Medvedev and Kanwal, 2008). In this case, evidence for the gamma-band activity being a different component from the broadband evoked-related potentials relies on the statistical independence between low- and high-frequency power in the LFPs. For this analysis the *full* window was used (see **Fig. 6A**). The power distributions of gamma (either 25-45 or 45-60 Hz) and low frequencies typically did not differ in our dataset (see **Fig. 7A, E** for a representative penetration). A systematic population analysis was performed to quantify the percentage of penetrations in the data for which there was evidence of statistical independence between the power of gamma and that of low-frequency potentials. As depicted in **Fig. 7B, F**, power distributions of gamma (in the 25-45 and 45-60 Hz ranges) and low frequencies (0-15 Hz) were significantly different from each other (2-sample Kolmogorov Smirnov tests, significance when p < 0.01) only in a small proportion (< 15%) of the total amount of penetrations, either in FAF or AC. This could suggest that gamma-band activity in these frequency ranges cannot be readily disentangled from a broadband power increase related to an onset response, assuming that if they were separable processes (i.e. gamma and low frequency activities) their power distributions would differ significantly (Medvedev and Kanwal, 2008).

**Fig. 7.**
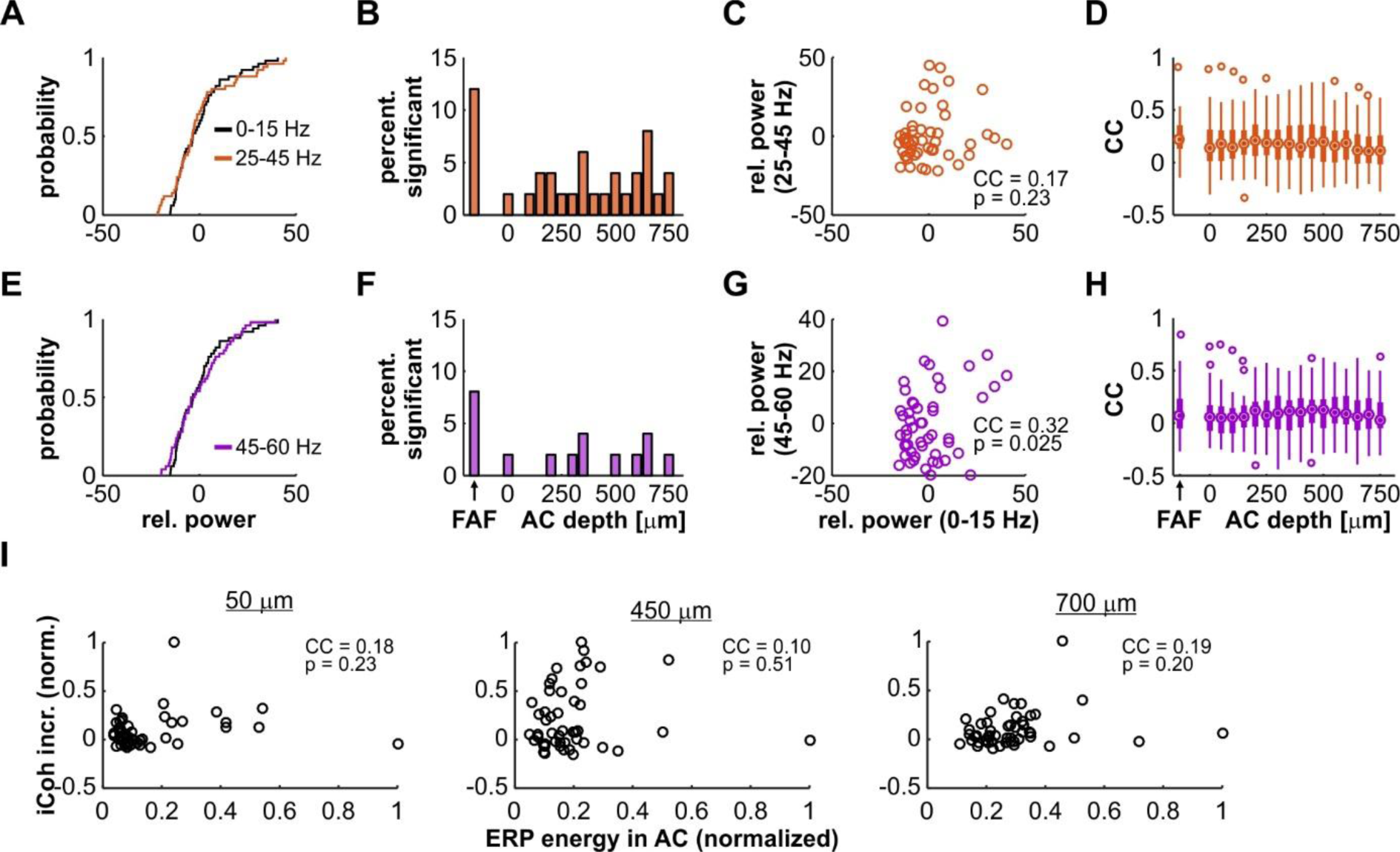
Power distributions of gamma-band and low-frequency LFPs in AC and FAF. (**A**) Distributions of the relative power of low-frequency (0-15 Hz; black) and gamma-band activity (25-45 Hz; orange) across trials, recorded from a single representative penetration in the FAF. These distributions were not significantly different from each other (2-sample Kolmogorov-Smirnov test, p = 0.84). (**B**) Percentage from the total number of penetrations (n = 50) for which the distributions of low-frequency and gamma (25-45 Hz) power were significantly different from each other at an alpha of 0.01, in FAF and at different depths of the AC. (**C**) Scatter plot and correlation coefficient (CC) of the trial-by-trial relationship between low-frequency and gamma band (25-45 Hz) power (n = 50 trials), for the same representative penetration shown in **A.** The CC was of 0.17, and it was not significant: p = 0.23. (**D**) Distribution, in FAF and at all AC depths, of CCs between gamma-band (25-45 Hz) and low-frequency power. The median in the AC across depths was of 0.17, whereas the median in the FAF was of 0.22. (**E-H**) Similar to **A-D**, but the gamma range considered was of 45-60 Hz (signaled in purple). In panel **E**, both distributions were also not significant from each other (2-sample Kolmogorov-Smirnov test, p = 0.84). The CC in **G** was of 0.32, and it was not significant at an alpha of 0.01 (p = 0.025). In panel **H**, the median across depths in the AC was of 0.08, whereas the median CC in the FAF was of 0.07. (**I**) Correlations between evoked-potential (ERP) energy in AC and gamma-band coherence increase (same as in Fig. 5) for three representative depths in the AC (at 50, 450 and **B.** 700 μm; all depths are shown in Fig. S3). Values are normalized for clarity. There were no significant correlations at any of the depths shown (p >= 0.2).

**Fig. 8.**
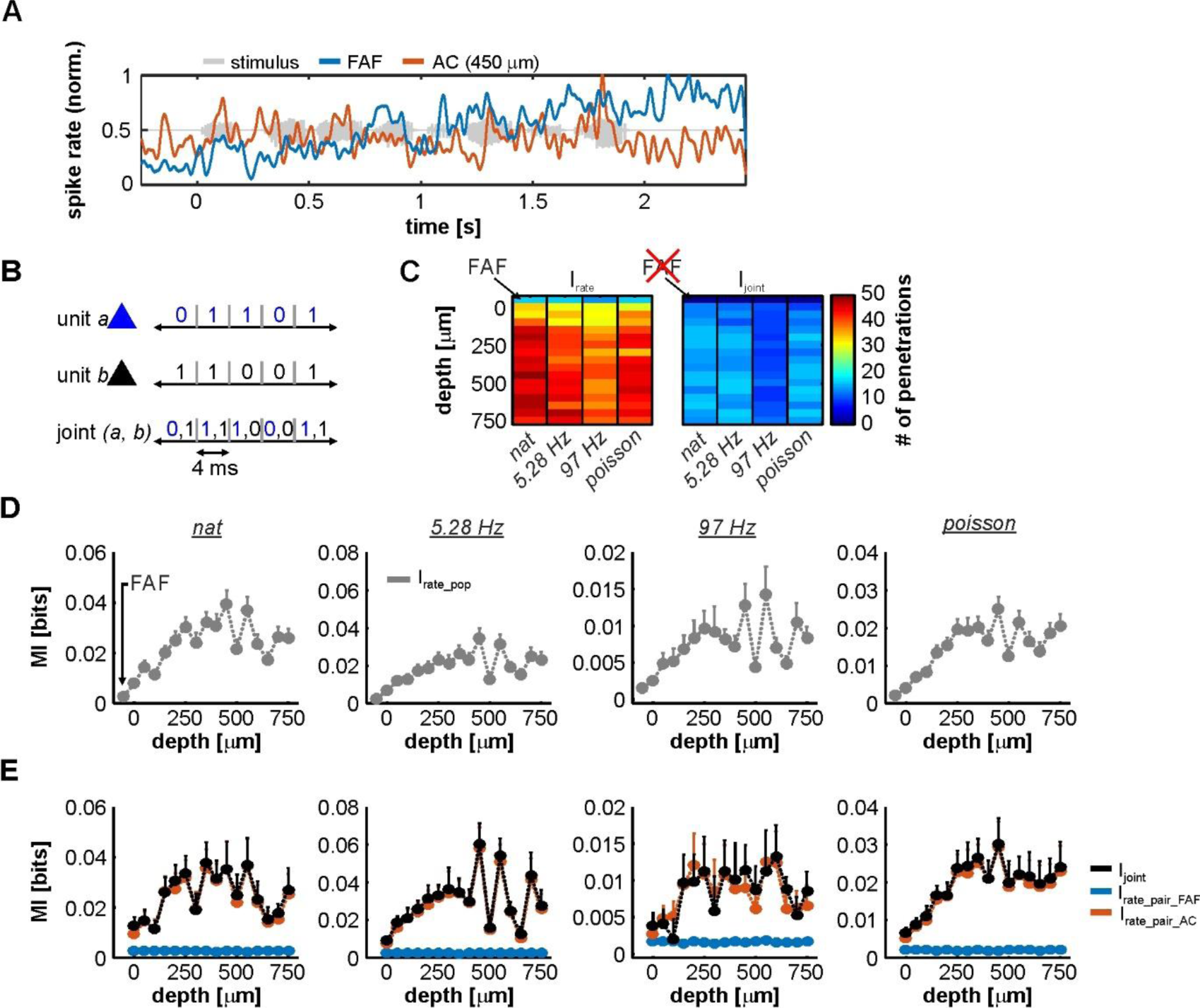
Information in rate codes of the FAF-AC circuit. (**A**) Spiking activity in an exemplary simultaneous recording from the FAF (blue) and the AC (at 450 μm, red). Note the difference in the response patterns between the two structures. (**B**) Schematic representation of the joint code used for information theoretic calculations. (**C**) Number of units used to calculate I_rate_ (left heatmap; 1 unit per penetration), for each stimulus (natural call, syllable train at 5.28 Hz, syllable train at 97 Hz, and Poisson syllable train) and depth in the AC (note that the first row in the heatmap corresponds to the FAF). Only units that had at least 0.1 bit/s of information were considered. The right heatmap depicts similar information, showing the number of pairs (FAF-AC spiking; 1 pair per penetration) used to calculate I_rate_ per stimulus and AC depth. In this case, FAF is missing because it is already part of each a pair. (**D**) Population I_rate_ in the FAF and across AC depths (note that the leftmost values correspond to the FAF), for each of the four stimuli presented (left to right). (**E**) I_joint_ (black) shown together with I_rate_ from AC (orange) and FAF (blue) units that conformed the pairs, across stimuli (i.e. I_rate_pair_AC_ and I_rate_pair_FAF_). There were no consistent significant differences between I_joint_ and I_rate_, when the latter was calculated for AC units (Wilcoxon signed rank tests, p > 0.05).

We reasoned, however, that a lack of significant differences between the distributions does not necessarily imply that the relative powers of gamma and low-frequencies are well correlated when considering trial specific information. Strong correlations would occur if low and high frequency relative powers were tightly determined by the strength of the broadband activation. Therefore, given a weak correlation, it could be argued that the dynamics of low- and high-frequency (gamma) LFPs might be more complex than an unspecific power increase related to the evoked response. We observed poor correlations, across penetrations in FAF and AC, between low-frequency and gamma-band relative power on a trial-by-trial basis (see, for example, **Fig. 7C, G**). The distribution of correlation coefficients for the population data is depicted in **Fig. 7D, H**. Overall, correlation coefficients were low, having a median in the FAF of 0.22 (25^th^ and 75^th^ percentiles: 0.12 and 0.35) for the 25-45 Hz gamma range, and of 0.07 (25^th^ and 75^th^ percentiles: −0.05 and 0.23) for the 45-60 Hz band. In the AC, the median across channels was of 0.17 (25^th^ and 75^th^ percentiles: 0.03 and 0.31) for the 25-45 Hz gamma, and of 0.08 (25^th^ and 75^th^ percentiles: −0.04 and 0.1) for the band of 45-60 Hz. Typically, no more than 20-25% of the penetrations in AC and FAF showed a significant correlation (significance when p < 0.01) between relative power at low-frequencies and gamma (25-45 Hz, median 20% of sites; 45-60 Hz, median 8% of sites; see **Fig. S2B**).

We also quantified how the overall energy of early activation correlates with the gamma coherence increase in the FAF-AC circuit. Gamma-band coherence increase (compared to spontaneous activity, see **Fig. 5**) was poorly correlated with evoked potential energy for all AC channels (median across channels: 0.15; 25^th^ and 75^th^ percentiles: 0.08 and 0.22), as illustrated in **Fig. 7I** for representative depths in AC, and in **Supplementary Fig. S3** for all depths. Although gamma-band activity is not straightforwardly separable from a broadband activation pattern, the data shown in **Fig. S3** and the poor trial-by-trial correlation between gamma and low-frequency powers (see above) suggest the possibility of interesting gamma-band dynamics in *C. perspicillata*’s FAF and AC.

### 2.5 Mutual information in FAF and AC spiking

We investigated spike-spike interactions in the FAF-AC network within an information theoretical framework (Shannon, 2001). Mutual information (MI or “information” throughout the text) between the stimuli and neuronal responses allows to quantify the theoretical ability of a neuron (or a set thereof) to represent the acoustic input on a single trial basis (see Methods). MI captures all non-linear dependencies of any statistical order in the data, and its quantification depends on the neural code being considered (Kayser et al., 2009). Here, we aimed to determine the coding abilities in AC, FAF, and a joint response from both structures based on a spike rate code (I_rate_ for single units, I_joint_ considering responses from AC and FAF together; (Kayser et al., 2009; Garcia-Rosales et al., 2018a)). A rate code was considered so that our results could be comparable with previous data obtained from *C. perspicillata*’s AC (Garcia-Rosales et al., 2018a).

Overall, we relied on an information theoretic approach because we observed that representations in AC and FAF were quite different and on occasions, at least in appearance, complementary. For example, **Fig. 6A** depicts spiking from two simultaneously recorded FAF and AC units in response to the natural stimulus. Note how the firing rate of the FAF unit increases as the stimulus progresses, whereas the AC unit is time-locked to slow temporal modulations in the stimulus and does not respond like its FAF counterpart. Information theory would allow to measure possible interactions between these responses in a quantitative manner.

**Figure 6B** shows a schematic of the rate code used to quantify MI for single units and joint responses. For subsequent analyses and in order to guarantee that all units considered were auditory-responsive (both in FAF and AC), we used only responses that provided at least 0.1 bit/s of information (with a window size of 4 ms this corresponds to 4×10^-3^ bits; see Methods). The number of units used to quantify I_rate_, per channel (including the FAF electrode, and across the AC linear probe) is depicted in **Fig. 6C**, left. Note that the number of penetrations is equivalent to the number of units per channel in this case: n ≥ 13 in FAF, and n ≥ 29 in AC. For paired responses (FAF-AC) the number of units considered was less because the inclusion criterion (I_rate_ ≥ 0.1 bit/s) had to be fulfilled by units in AC and FAF simultaneously (n ≥ 8 pairs; **Fig. 6C, right**).

Population values of I_rate_ for the FAF and the AC at different depths are depicted in **Fig. 6D**, for each stimulus. Two main conclusions can be drafted from this figure. (i) In the AC, the highest information about the stimuli was found at depths between 200-650 μm. (ii) Neurons in the FAF were on average less informative than AC ones, but were well above the limit set in the inclusion criterion across stimuli (*nat*: 0.56 ± 0.09 bits/s, 5.28 Hz train: 0.45 ± 0.06 bits/s, 97 Hz train: 0.48 ± 0.04 bits/s, Poisson train: 0.42 ± 0.05 bits/s; given as mean ± s.e.m). We also quantified the information provided by paired neuronal responses from both regions (I_joint_), which is illustrated as black traces in **Fig. 6E**. This figure also shows a direct comparison between I_joint_ and the I_rate_ of FAF and AC neurons that conform each pair (I_rate_pair_FAF_ and I_rate_pair_AC_, respectively). The contribution of I_rate_ from the FAF was significantly smaller than the contribution of I_rate_ from the AC (FDR-corrected Wilcoxon signed rank tests, p_corr_ < 0.05; corrected p values of all comparisons are given in **Fig. S4**). Although I_rate_ in the FAF was always > 0.1 bit/s, we did not observe I_joint_ to be significantly higher than the I_rate_ from the AC (I_rate_AC_; orange traces in the **Fig. 6E**), across electrodes and stimuli tested (**Fig. S7**). We did observe I_joint_ to be well correlated with the sum of I_rate_ calculated using the information of FAF and AC spiking (I_sum_ = I_rate_pair_FAF_ + I_rate_pair_AC_; **Figs. S5-S9**).

When considering the interactions of spiking across structures in terms of the codes defined in this study, the linear relationship between I_sum_ and I_joint_, evident in **Figs. S5-S9**, would support the notion of “independence” in the information provided by the spiking in each structure. Independence arises theoretically when I_joint_ = I_sum_, and implies that each structure represents different aspects of the sensory stimulus. However, the results illustrated in **Fig. 6E** suggest that independence cannot be inferred from the data with certainty. Mathematically, independence would require I_joint_ to be higher than both I_rate_ in AC and FAF simultaneously, which was not fulfilled at a population level: I_joint_ was not significantly different than I_rate_ obtained from AC units (see **Fig. S4**). Calculating information estimates using time windows of up to 12 ms yielded comparable results, indicating that the temporal resolution of the codes had little effect in the described outcomes (data not shown). Overall, our quantification of stimulus-related spiking information, occurring simultaneously in the AC and FAF, suggests different sound coding strategies in these two structures. These observations are further addressed in the Discussion section.

## 3 Discussion

In this study we investigated the functional connectivity between frontal and auditory cortical regions of the bat *Carollia perspicillata*. Specifically, we examined the coupling dynamics of the FAF, a frontal area which receives auditory afferents from cortical and subcortical structures, and the primary AC. Functional connectivity was assessed during spontaneous activity and during the processing of natural and artificially generated acoustic sequences. Our main results are: i) LFPs recorded simultaneously in both regions suggest that the FAF receives faster auditory inputs relative to the AC, yet these inputs do not necessarily elicit faster spiking in the FAF; ii) during spontaneous activity, the FAF-AC network is coupled in low frequencies (up to 12 Hz), with stronger coherence values at deep layers of the AC; iii) while acoustic stimulation does not considerably alter the default low-frequency coupling, auditory-evoked gamma-band synchronization in the FAF-AC circuit emerges upon sound presentation; iv) considering a spiking rate code, the FAF is less informative than the AC about the acoustic stimuli, while a joint code using simultaneous spiking from both regions suggests that FAF and AC engage in distinct coding dynamics. Altogether, our data shed light onto how distant brain areas in the mammalian brain engage in sound representation. The results of this paper are summarized in **Fig. 9**.

**Fig. 9.**
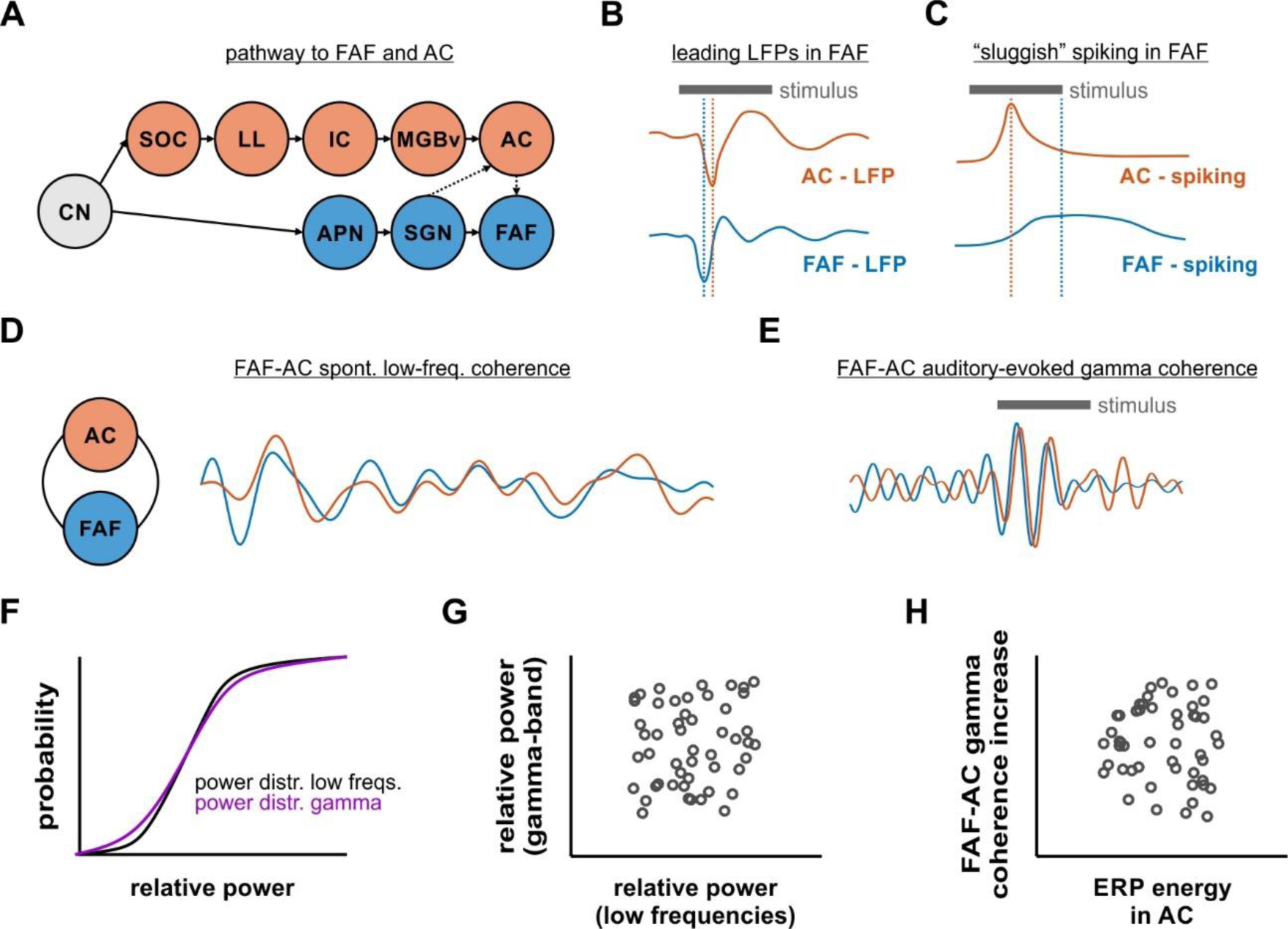
Functional coupling dynamics in the FAF-AC circuit of *C. perspicillata*. (**A**) Representation of the auditory pathway to the FAF and the AC (CN, cochlear nucleus; SOC, superior olivary complex; LL, lateral lemniscus; IC, inferior colliculus; MGBv, ventral division of the medial geniculate body of the thalamus; AC, auditory cortex; APN, anterolateral periolivary nucleus; SGN, suprageniculate nucleus of the thalamus; FAF, frontal auditory field). (**B**) Schematic representation illustrating that stimulus-related LFPs in FAF lead relative to those in the AC (see Fig. 1). (**C**) Fast LFP responses in FAF do not necessarily elicit fast spiking responses (schematic). Neurons in the frontal region typically respond more “sluggishly” than their auditory cortical counterparts. (**D**) FAF and AC were coherent in low-frequencies during spontaneous activity (i.e. in the absence of sound stimulation). (**E**) During acoustic processing in passive listening animals, low-frequency coherence was unaltered in the FAF-AC circuit, although there was an emergence of auditory-evoked gamma band coherence in the network. Traces in panels D and E are based on data shown in Figs. 3 and 4. Note that, for illustrative purposes, the temporal scales and amplitudes in **D** and **E** are not comparable. (**F**) Distributions of power in low and gamma-band frequencies were typically not significantly different from each other. However, there was very weak trial-by-trial correlation between low-frequency and gamma power (**G**), as well as very low correlations between event-related potential (ERP) energy and gamma coherence increase across penetrations (**H**).

### 3.1 Auditory afferents into the FAF

Stimulus-related LFPs recorded simultaneously from FAF and primary AC, at various depths in the latter structure, indicate the presence of fast synaptic inputs into the frontal region that precede those arriving into the AC even at input layers. The work of Kobler, Casseday, and colleagues in the late 1980s (Kobler et al., 1987; Casseday et al., 1989), showed that the FAF receives auditory afferents via a non-canonical pathway that bypasses major auditory centres in the midbrain, including the inferior colliculus (IC). In this pathway, acoustic information from neurons in the cochlear nucleus is sequentially relayed to the anterolateral olivary complex, the suprageniculate nucleus of the thalamus (SGN), and from there into the FAF. Thus, although auditory inputs reach the frontal region also through the AC, acoustic information may reach the frontal field, directly from the cochlea, in as few as four synapses (Kobler et al., 1987). Synaptic currents related to inputs from the SGN into the FAF could lead to changes in LFPs (Buzsaki et al., 2012) faster than their counterparts at input layers of the AC, which are predominantly driven by thalamocortical synapses originating in the ventral region of the medial geniculate body. Therefore, a rapid, non-canonical pathway into FAF accounts for our observations regarding the temporal relation between LFPs in both regions studied.

The results described in this manuscript indicate that, although sound-related LFPs from the FAF lead those from the AC, the neuronal spiking latencies are shorter in AC. In other words, our data indicate that faster inputs in FAF are not sufficient to elicit faster spiking in most cases.

Electrophysiological studies in frontal auditory areas have shown that neuronal responses are usually of relatively large latencies (although short latencies are also to be found), sparse and of high variability (Newman and Lindsley, 1976; Eiermann and Esser, 2000; Kanwal et al., 2000; Plakke and Romanski, 2014). Recent data from *C. perspicillata*’s FAF highlighted the possibility that the sparseness in the response properties of frontal neurons could be explained by slow, low-threshold, and long-lasting synaptic dynamics, at least considering projections from the AC (Lopez-Jury et al., 2019). These slow, long-lasting synaptic dynamics could support sensory integration, by conditioning FAF neurons to spike after accumulating synaptic inputs over time, and/or to integrate multiple synaptic inputs originating from different sensory modalities. Certainly, cross-modal sensory integration occurs in the frontal cortex (Fuster et al., 2000; Romanski, 2007; Hwang and Romanski, 2015), whereas integration over relatively long timescales appears to be a feature of higher-order cortical areas in general (Runyan et al., 2017).

Fast acoustic afferents, even without eliciting reliable stimulus-evoked spiking, imply nonetheless that auditory information is already present in the FAF before it receives inputs from the AC. Within the predictive coding framework (Arnal and Giraud, 2012; Friston, 2018), it is then possible to hypothesize that the frontal field may be a region where prediction errors could also be generated.

That is, the rapid non-lemniscal pathway could relay faithful information into FAF about the auditory stimuli, which can in turn be compared with information received from primary AC and the thalamus. Alternatively, a further proposition would be that prediction errors, relayed “upwards” along the auditory hierarchy (Carbajal and Malmierca, 2018), and generated in the SGN and the AC, are integrated in the FAF. The result of such integration could in turn be used to update the “expectations” of the system. In rodents, prediction error signals appear all along the auditory pathway, but occur more strongly in higher-order structures (Parras et al., 2017), while prediction error signals related to sounds have also been described in the frontal cortex of rats (Imada et al., 2012) and humans (Durschmid et al., 2016; Durschmid et al., 2018). The involvement of the FAF in predictive coding could be thoroughly tested in future experimental work.

A number of FAF neurons project directly to the superior colliculus (SC), a structure related to motor control of head and pinna movement in bats (Kobler et al., 1987). The presence of early acoustic information in FAF, the fact that frontal neurons in *C. perspicillata* show preference for naturalistic echolocation acoustic stimuli (high-frequency pulses used to navigate by bats; (Eiermann and Esser, 2000)), and the existence of projections from FAF into motor-related structures such as the SC, suggest that the FAF plays an important role in coordinating auditory-guided behaviour. This would be in line with proposed roles of prefrontal cortex in motor control (Risterucci et al., 2003), including volitional motor (vocal) production after acoustic stimulation (Hage and Nieder, 2015). However, whether neuronal activity in the FAF mediates motor outputs is still to be tested in depth.

### 3.2 FAF-AC synchronization during spontaneous activity

We examined the default (i.e. in the absence of external stimulation) functional connectivity in the FAF-AC network in terms of oscillatory coherence (**Fig. 3**). The data presented in this manuscript show that simultaneously recorded LFPs from both structures are phase-synchronized in low-frequencies of the spectrum (1-12 Hz), although more strongly so in the delta band (1-4 Hz).

Empirical evidence points towards a role of oscillatory coherence in the coupling of distant brain regions, a view that is summarized in proposed theoretical mechanisms such as the communication-through-coherence framework (Fries, 2015). Several studies have pinpointed an involvement of low-frequency synchronization in the functional coupling between the frontal cortex and a variety of brain structures. Coherent oscillations between distant cortical areas (including the frontal cortex) in the low-frequency range correlate with working memory (Daume et al., 2017), fear memory consolidation (Popa et al., 2010), attentional selection (Womelsdorf and Everling, 2015), and long-term fear recall (Cambiaghi et al., 2016; Karalis et al., 2016). In the auditory domain, top-down control exerted from frontal cortical areas, through low-frequency oscillatory activity, increases coupling to speech signals in the human AC (Park et al., 2015). Our results indicate that, in the bat brain, frontal areas that participate in audition are functionally interconnected by means of low-frequency LFPs with primary auditory cortex. Critically, such coupling does not require external input, which hints towards the presence of a default synchrony in the fronto-auditory cortical circuitry. The latter could constitute a functional basis for high-order, interareal auditory processing in the mammalian brain.

Because recordings in the AC were performed with a laminar probe, we were able to study the laminar dependence of FAF-AC synchrony in AC. During spontaneous activity, low-frequency coherence was strongest in deep layers (depth > 700 μm), but the strength of coherence was only affected by depth in the delta band. Interestingly, a recent study revealed that, also during spontaneous activity, spike-LFP synchronization was strongest in deep layers of the AC (Garcia-Rosales et al., 2019). Such spike-LFP coupling was associated to the presence of discrete spontaneous states of increased spiking rate (UP-states) in laminae V and VI. The origins of deep layer UP-states are unclear, but it has been hypothesized that they could be driven by higher-order structures (Sakata and Harris, 2009). In the current study, we observed a putative higher-order auditory structure (the FAF in the frontal lobe) synchronized via a low-frequency oscillatory channel with deep layers of a primary sensory area (the AC). From the phase correlation of delta-band LFPs in the FAF-AC network, it is possible to speculate that delta oscillations in FAF could modulate UP-states in AC. However, we note that causality cannot be inferred from our current dataset and needs to be addressed thoroughly with further experimental approaches.

In all, the “default” coupling in the FAF-AC circuit is supported by the presence of anatomical connections between frontal and auditory cortices (Kobler et al., 1987)). Although it has to be properly addressed, we propose that homologous frontal regions tasked with audition in other species may be functionally interconnected with AC in a similar manner. This possibility is still unexplored, yet addressing this question might be crucial for unravelling the mechanisms of high-order auditory processing, cognition, and behaviour based on audition.

### 3.3 Functional coupling in the FAF-AC network during acoustic processing

Functional coupling between FAF and AC was also addressed in the context of acoustic processing (**Figs. 4, 5**). Animals were exposed to four distinct acoustic streams, including a conspecific distress vocalization, and three “trains” constructed by repeating a distress syllable at distinct rates.

Independently of the stimulus used, we observed little change in the low-frequency phase synchrony of the FAF-AC network, as compared to spontaneous activity. This is a puzzling result which, assuming that low-frequency oscillatory coupling in the network is useful for auditory perception, could in principle be explained largely by our experimental design. Although the animals were awake during the experiments, they listened to the sequences passively: i.e. they were not expected to behave in response to the stimulus, and other variables (e.g. attentional processes) were not modulated according to a controlled experimental approach. Thus, statistically negligible and unreliable changes in the low-frequency dynamics of FAF-AC connectivity could be explained by the fact that the passive listening of acoustic streams is not sufficient to alter the default functional coupling in the network. Whether attention (a top-down process) or behavioural planning could modify the neuronal connectivity in the circuit by either enhancing it or decreasing it, as compared to spontaneous activity, needs to be tested in further research.

Experimental evidence suggests that oscillatory activity in the gamma range is crucial for neuronal computations, including sensory processing and cognitive mechanisms (Fries, 2009). Previous studies have shown the presence of gamma-band activity in primary auditory cortex of rats (Vianney-Rodrigues et al., 2011), monkeys (Brosch et al., 2002) and bats (Medvedev and Kanwal, 2008) during passive listening. These studies reported the presence of gamma oscillations at even later time periods (150-300 ms) in what could be considered “induced” (as opposed to “evoked”) activity, not-locked to the auditory stimulus. Our results show that, past 90 ms and up to 180 ms after stimulus presentation, such late, non-locked gamma oscillations are relatively scarce (∼5% of out of 50 penetrations). These results should not be taken as evidence for the lack of late gamma oscillations in the AC or FAF of *C. perspicillata*, because later periods (190-300 ms after sound presentation) in which oscillatory activity may have occurred were not analysed due to the nature of the stimulus (note that the second syllable presentation of the 5.28 Hz occurs already at ∼189.4 ms). Further research can be aimed at detecting gamma activity at later time points in the cortex of *C. perspicillata*.

In humans, sources of auditory-evoked gamma-band activity (aeGBA) can be found both in primary auditory and frontal (anterior cingulate cortex, ACC) cortices (Mulert et al., 2007; Polomac et al., 2015). Frontal aeGBA is modulated by attention and correlates with performance in auditory detection tasks (Debener et al., 2003; Gurtubay et al., 2004). Moreover, aeGBA in the frontal lobe and gamma-band synchronization between frontal and auditory cortical regions correlate too with task difficulty (Mulert et al., 2007; Polomac et al., 2015), suggesting a role of gamma-band coherence in a fronto-auditory cortical circuit for cognitive control in audition. In fact, disorders of the central nervous system such as schizophrenia are marked by a dysregulation of aeGBA in frontal regions (Cho et al., 2006; Leicht et al., 2010; Curic et al., 2019), further supporting the importance of gamma-band activity for cognition.

Our data indicate that low-gamma (25-45 Hz) coherence in the FAF-AC circuit significantly increases with sound presentation, independently of the stimulus considered. This supports a role of gamma synchronization between frontal and auditory cortices for auditory processing, although the functional significance of aeGBA and its coherence across cortical areas should be considered with care. Coherent activity does not imply directly that there exists effective communication between two given regions. Moreover, the nature of the gamma activity is also to be examined cautiously. Here we attempted to disentangle gamma oscillations from a frequency unspecific power surge related to auditory evoked responses (**Fig. 7**), which would in principle explain gamma coherence between FAF and AC. Based on a method proposed by Medvedev and Kanwal (2008), we did not observe statistical evidence supporting that the distributions of low-frequency and gamma-band LFP power were significantly different from one another across penetrations in FAF and AC (**Fig. 7B, F**).

However, we did observe, across multiple penetrations, a lack of trial-by-trial correlation between the LFP power in the abovementioned frequency bands (**Fig. 7C, D, G, H**, and **Fig. S2B**). In addition, our data also showed a very weak correlation between the energy of the evoked response in the LFP and the gamma-band coherence increase (**Fig. 7I** and Fig. S3). While these results do not conclusively demonstrate that gamma-band activity can be separated from a frequency unspecific onset response, they hint towards the possibility of evoked gamma activity being an important component for audition, as suggested by previous work (Brosch et al., 2002; Medvedev and Kanwal, 2008; Vianney-Rodrigues et al., 2011). The functional roles of onset-related gamma activity in the FAF-AC circuit of *C. perspicillata* (and in the mammalian auditory system in general) should be thoroughly addressed with dedicated experimental approaches in the future.

We would like to note that it remains speculative what might be potentially signalled by the FAF-AC gamma synchronization reported in this study. Short onset-related coherence increase might convey information about the presence of an acoustic stimulus, but not necessarily allow to elucidate the stimulus’ spectrotemporal features. In the AC of the bat *Pteronotus parnelii*, Medvedev and Kanwal (2008) reported that the spectral properties of the gamma component of the response could be used to differentiate among a battery of conspecific communication calls. In primary visual cortex, for example, distinct characteristics of stimulus-induced gamma rhythms (e.g. peak frequency or amplitude) encode for distinct properties of presented visual stimuli (e.g. contrast or orientation; (Hermes et al., 2015; Murty et al., 2018)), although it has been argued that the variability of gamma based on stimulus properties may constrain the utility of the rhythm for complex integrative computations, at least in early visual cortex (Henrie and Shapley, 2005; Ray and Maunsell, 2010; Bartoli et al., 2019). Our stimulus set is not ideal to determine in an unbiased manner whether the nature of FAF-AC gamma synchronization changes given the spectrotemporal characteristics of the stimuli, in particular because LFPs synchronize to a stimulus’ temporal structure also in the gamma range (see (Hechavarria et al., 2016b; Garcia-Rosales et al., 2018a). The latter could alter the spectral patterns of coherence without necessarily meaning that the nature of the underlying coherence is changing, which is an artefact that needs to be controlled for. The careful and systematic variation of acoustic properties of sounds could be used in further research to explore in full the patterns of FAF-AC gamma-band coherence and its role for audition.

### 3.4 Mutual information in the FAF-AC circuit

Quantifying the amount of information provided by FAF and AC about the acoustic stimuli revealed that the former structure was, in comparison, significantly less informative than the latter. In addition, when considering responses from frontal or auditory cortices in a joint rate code, it was not possible to determine a clear population trend towards independence, redundancy or synergy between the spiking activities of both structures, from an information theoretic perspective. We propose two candidate explanations for our results, which need not be mutually exclusive. First, it is possible that the way in which FAF encodes incoming auditory stimuli is not sufficiently well-captured by means of a code based on spiking rate, and therefore the true encoding capabilities could be underestimated by assuming such scheme (see (Masquelier, 2013; Insanally et al., 2019)). Second, we note that the fact that the FAF conveys, in comparison, significantly less information than the AC, is an expected result as the AC is a structure specialized for auditory computations. As part of the frontal cortex, it is plausible that the FAF encodes for other variables that go beyond acoustic features (e.g. ethological relevance of the sound, multimodal sensory information, etc.). In that case, FAF units would yield low I_rate_ values when attempting to quantify their abilities to encode a sound based on relatively simple methodological approaches, which rely solely on acoustic processing. The former allows to hypothesize that FAF and AC might engage in distinct, non-overlapping coding strategies.

## 4 Methods

### 4.1 Animal preparation and surgical procedures

All experimental procedures were performed in compliance with current German and European regulations on animal experimentation. Experiments were approved by the Regierungspräsidium Darmstadt (experimental permit #FU-1126). The study was performed on 5 adult bats of the species *Carollia perspicillata* (all males). Animals were obtained from a colony in the Institute for Cell Biology and Neuroscience, Goethe University, Frankfurt am Main. Bats used for experiments were kept separately from the main colony.

Before undergoing surgical procedures, bats were anaesthetized with a mixture of ketamine (10 mg*kg^-1^, Ketavet, Pfizer) and xylazine (38 mg*kg^-1^, Rompun, Bayer). For surgery and any subsequent handling of the wounds, local anaesthesia (ropivacaine hydrochloride, 2 mg/ml, Fresenius Kabi, Germany) was applied subcutaneously in the scalp area. A rostro-caudal midline incision was made in the scalp, after which skin and muscle tissues were removed carefully in order to expose the skull. A sufficiently large area of the bone was also exposed to make possible the attachment of a custom-made metal rode (1 cm length, 0.1 cm diameter), used during recordings to fixate the animal’s head. The rod was attached with dental cement (Paladur, Heraeus Kulzer GmbH, Germany). Animals were given at least one full day of recovery after surgery, and before experiments were performed upon them. The AC and the FAF were located based on well-establish landmarks such as blood vessel patterns, and the sulcus anterior (see (Esser and Eiermann, 1999; Eiermann and Esser, 2000)). On the first day of recordings each cortical region was exposed by cutting a small hole (∼ 1 mm^2^) in the skull with a scalpel blade.

Recordings, which lasted no more than 4 hours a day, were performed chronically on awake bats. Water was given to the animals at a period of approximately 1-1.5 hours. Between recording sessions, a bat was allowed to recover for at least a full day. Experiments for the day were halted if the bat showed any sign of discomfort.

### 4.2 Electrophysiological recordings

Recordings were made inside an electrically isolated and sound-proofed chamber. Inside the chamber, bats were placed upon a custom-made holder which was kept at a constant temperature of 30 °C with a heating blanket (Harvard, Homeothermic blanket control unit). A speaker (NeoCD 1.0 Ribbon Tweeter; Fountek Electronics, China), located inside of the chamber 12 cm away from the animal’s right ear (contralateral to the hemisphere were recordings were performed), was used for free-field stimulation. The speaker was calibrated using a ¼-inch microphone (Brüel & Kjær, model 4135, Denmark), which was connected to a custom-made amplifier.

Data were acquired from the bat’s left AC as described in a previous study (Garcia-Rosales et al., 2019). Neurophysiological data were recorded from the AC using 16-channel laminar electrodes (Model A1×16, NeuroNexus, MI; impedance: 0.5–3 MΩ), with a channel separation of 50 μm. The probe was carefully inserted into the brain perpendicular to the cortical surface using a piezo manipulator (PM-101, Science 455 products GmbH, Hofheim, Germany) until the top-channel was barely visible on the surface of the tissue. Thus, we were able to record from depths ranging 0-750 μm, reaching all layers in AC. Histological confirmation of the extent of the electrodes inside the cortex are detailed elsewhere (Garcia-Rosales et al., 2019). Recordings were made in primary AC, although we cannot discard the presence of columns from high frequency fields (Esser and Eiermann, 1999). The laminar probes were connected to a micro-amplifier (MPA 16, Multichannel Systems MCS GmbH, Reutlingen, Germany), and acquisition was done via a portable multichannel system with integrated analogue-to-digital converter (Multi Channel Systems MCS GmbH, model ME32 System, Germany) with a sampling frequency 20 kHz and a precision of 16 bits. Data acquisition was on-line monitored and stored in a computer using MC_Rack_Software (Multi Channel Systems MCS GmbH, Reutlingen, Germany; version 4.6.2).

For recordings in the FAF, a single carbon electrode (Carbostar-1, Kation scientific; Impedance at 1 kHz: 0.4–1.2 MΩ) was inserted into the frontal region of the left hemisphere and lowered to depths of ∼300-450 μm with the aid of a second piezo manipulator (same characteristics as the previous one). The electrode was connected to a micro-amplifier which was also connected to the integrated multichannel recording system as described above. It was possible to use the same hardware because the integrated system accommodates up to 32 simultaneous channel recordings. Ground and reference electrodes (silver wires) were inserted as to only touch the dura mater of non-auditory regions of the bat’s brain, preferentially located in occipital areas of the contralateral hemisphere.

### 4.3 Acoustic stimulation

Acoustic stimulation was controlled from the recording computer using a custom-written Matlab (version 7.9.0.529 (R2009b), MathWorks, Natick, MA) software. As acoustic stimuli we used a natural distress call from *C. perspicillata* and three synthetic trains constructed from a single distress syllable, repeated at different rates. Procedures for recording the natural sequence are described in a previous study (Hechavarria et al., 2016a). The call is representative of *C. perspicillata*’s vocal repertoire, and has been used by us in previous studies addressing auditory processing at the level of the AC (Hechavarria et al., 2016b; Garcia-Rosales et al., 2018a; Garcia-Rosales et al., 2019). Distress calls of *C. perspicillata* exhibit two prominent, coexistent temporal modulations: the syllabic- and bout rhythmicities. Syllabic rates in *C. perspicillata’*s distress utterances are in the range of > 30 Hz (median, 71.4 Hz; iqr: 57.1 Hz), whereas bouts (groups of syllables emitted in close sequence) are repeated with rates typically < 12 Hz (Hechavarria et al., 2016a). In the natural call used here, syllables are repeated on average with a rate of 63.7 Hz (see (Garcia-Rosales et al., 2018a)), whereas bouts are uttered with a rate of 4 Hz (i.e. 8 bouts in 1.96 s).

To emulate the temporal dynamics of the communication sequences, a stereotypical distress syllable was used to construct artificial acoustic sequences. A first syllabic train had a repetition rate of 5.28 Hz, matching the slow temporal dynamics of *C. perspicillata*’s distress utterances. A second one, with a repetition rate of 97 Hz, was used to simulate fast temporal dynamics in communication streams. Finally, we constructed a syllabic train where syllables were repeated in a Poisson-like manner, with an average rate of 70 Hz. This simulated fast-repetition rates without any periodicity and without a slow temporal structure. The 5.28 and 97 Hz trains had a duration of 2 s, while the Poisson train was 4 s long. The syllables had an intensity of 70 dB SPL (root-mean square), close to the intensity of the natural call (see (Garcia-Rosales et al., 2018a)).

Sounds were digital-to-analogue converted by means of a sound card (M2Tech Hi-face DAC, 384 kHz, 32 bit) and amplified (Rotel power amplifier, model RB-1050) in order to be presented through the speaker inside of the chamber. Prior to presentation the call and syllabic trains were down-sampled to 192 kHz, and low-pass filtered (80 kHz cut-off). All sounds were pseudorandomly presented 50 times each, with and inter-stimulus interval of 1 s. A period of 300 ms, and another of 500 ms, was appended at the beginning and the end of each sequence, respectively.

Prior to any acoustic stimulation, per penetration, electrophysiological data were acquired for a period of 180 s. These data were used for coherence analyses during spontaneous activity.

### 4.4 Separation of spiking activity and local-field potentials

All analyses were performed offline with custom-written Matlab (version 8.6.0.267246 (R2015b)) scripts. Initially, the raw electrophysiological signal from each channel (all electrodes in AC and FAF) was bandpass filtered (fourth-order Butterworth filter) in order to extract traces pertaining spiking activity (300-3000 Hz cut-off frequencies) and LFPs (0.1-300 Hz cut-off frequencies). For computational reasons, LFP data were down-sampled to 1 kHz and stored for subsequent analyses.

Spike detection and sorting from FAF and AC electrodes were performed using the SpyKING CIRCUS toolbox (Yger et al., 2018). Spike detection threshold was set at 5 median absolute deviations from the noise baseline, and spike sorting was done automatically by the SpyKING CIRCUS algorithm based on the probe’s geometry to avoid detecting the same templates in two adjacent electrodes. Each template was assigned to the electrode where its amplitude was the strongest. Per electrode (either in AC or FAF), we chose as representative spiking the template with the highest spike count. Spiking responses relative to a single electrode are referred to as a “unit” in the manuscript.

### 4.5 Spike latency estimation

Spike latency was defined as the time point in which a unit’s spiking rate was statistically different from the expected rate during spontaneous activity, based on a previous study (Chase and Young, 2007). In brief, the algorithm proposed by Chase and Young compares a unit’s response to a stimulus across several time windows, with the expected spiking rate under the assumption that the unit fires spontaneously with Poisson statistics, given a certain rate. A unit’s firing rate in the 250 ms silence period before stimulus onset, across the 50 repetitions from all stimuli tested (a total 200 trials), was considered its spontaneous spiking rate for the abovementioned assumption. The response of a unit to a certain stimulus (i.e. spiking after stimulus onset) was pooled across trials. Taking this pooled response, the probability of observing at least *n* spikes in a given window *t_n_* (after stimulus onset), assuming Poisson firing in the absence of acoustic inputs, can be defined as follows (Chase and Young, 2007):

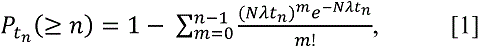

where N is the number of repetitions of the given stimulus, and *λ* is the spontaneous firing rate. Starting from stimulation onset, the probability that each elicited spike indicates a stronger than chance deviation in rate from the firing rate estimated in the absence of stimulation (the 250 ms window), is taken as the probability that the spontaneous firing rate would have produced that particular spike as the last of *n* spikes in a window *t_n_*. In this context, *t_n_* is the width of the window containing the *n* spikes observed so far. Hence, the time of the first spike for which the aforementioned probability is sufficiently low (here, *P_t__n_* (≥ *n*) < 10^−5^) is considered as the unit’s latency. This method circumvents caveats regarding classical peak latency estimations using peri-stimulus time histograms or spike-density functions over time (Levakova et al., 2015).

### 4.6 Interareal coherence analyses

All coherence analyses were done using the Chronux toolbox (Bokil et al., 2010). As a metric of interareal phase synchronization we used the imaginary part of the coherency (“iCoh” in the manuscript; (Nolte et al., 2004)), both during spontaneous activity and acoustic processing.

Coherency is complex value that measures phase consistency between two time series, across several trials. The coherency between two signals *x* and *y*, at a certain frequency ω, can be defined as follows (Bastos and Schoffelen, 2015):

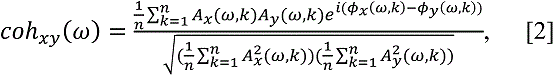

where *A_x_(ω, k)* and *A_y_(ω, k)* are the amplitudes of signals *x* and *y* at frequency ω and trial *k*, while *coh_xy_* represents the coherency between both signals, and *n* is the number of trials per stimulus (n = 50). Coherency is a complex quantity, but its absolute value ranges from 0 to 1, indicating the relative, normalized strength of phase synchronization between time series. Taking the imaginary part of *coh_xy_* is a straightforward manner to remove non phase-lagged interactions that could be attributable to, for example, passive field spread or common referencing (Bastos and Schoffelen, 2015). In order to minimize further common influences related to the temporal structure of the stimuli, we subtracted from each trial the mean (across all trials of a given stimulus) LFP of each channel (Kikuchi et al., 2017), and calculated coherence using the de-meaned traces. Note that this could affect low frequencies more than high frequencies, because the latter are more sensitive to temporal jitter. Additionally, while the approach alleviates obtaining simply stimulus-evoked coherence, it could also mask phase-locking that is time-locked to the stimulus but not entirely attributable to acoustic temporal features.

From the 180 s trace of spontaneous activity recorded simultaneously from FAF and AC penetrations, 50 chunks of 3 s length each were taken. The precise time at which chunks started was chosen randomly in a way that the resulting sub-segments would still be non-overlapping. Each of these paired chunks from FAF and AC were treated as a trial, and iCoh was estimated from all 50 of them for the corresponding penetration (data shown in **Fig. 3B**). A surrogate calculation was performed whereby the precise phase relationship between FAF and AC “trials” in spontaneous activity was abolished. This was accomplished by pairing FAF chunks with AC chunks randomly. The former affects the timing of the phase-relationships but maintains the overall power across selected chunks. The pairing was performed randomly a sufficiently large number of times (250), and iCoh was calculated at each repetition of the surrogate analysis. Thus, it was possible to obtain a distribution of iCoh values that represented coherence estimates in the absence of consistent phase relationships between AC and FAF during spontaneous activity. The iCoh calculated from the original data was then related to the surrogate iCoh values by means of z-normalization (z-iCoh). At a population level, lack of consistent phase coherence between FAF and AC at a certain frequency would yield z-iCoh values close to 0.

Time-frequency resolved iCoh values were obtained by means of coherogram calculations (*cohgram* function in Chronux; data depicted in **Figs. 3C, 4** and **5**). A time-resolved approach allowed us to examine changes of coherence over time while animals listened to acoustic streams. Each coherogram was constructed by calculating coherence in a sliding window of 200 ms length, which was advanced in steps of 2 ms. Because of the spectral resolution due to window length, the coherence spectrum for frequencies below 4 Hz could not be estimated with precision. As with any time-frequency resolved approach, there is a compromise between temporal and spectral resolutions, which we empirically found to be best balanced with a 200 ms window. All power spectra in the time-resolved analysis were obtained with the multitaper method (Percival and Walden, 1993), available in the Chronux toolbox, using 3 tapers and a time-bandwidth (TW) product 2.

The frequency range of 4-12 Hz was used as a representative of low-frequencies in the spectrum, and the 25-45 Hz band was considered as low-gamma. For comparing iCoh values during sound presentation vs. iCoh values during spontaneous activity, we calculated time-resolved iCoh during spontaneous activity using the same segments with which non-time resolved coherency was calculated (shown in **Fig. 3C**). Because the length of the spontaneous segments (3 s) was not precisely equal to the length of the stimuli (the Poisson process, for example, was 4 s long), we collapsed the time-resolved spontaneous iCoh values in the temporal dimension (median across timepoints per frequency). Thus, it was possible estimate the percentage increase during sound processing in a time-resolved manner as follows:

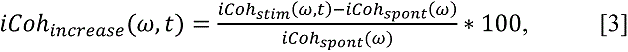

where *iCoh_stim_(ω, t)* is the iCoh value during stimulus presentation at frequency ω ant time *t*, while *iCoh_spont_(ω)* is the collapsed time-resolved iCoh during spontaneous activity at the same frequency. The percentage increases were narrowed to the frequency bands of interest (i.e. 4-12 and 25-45 Hz) by calculating the median iCoh increase in the corresponding frequency range over time (data depicted in **Fig. 5A, D, G, J**). For evaluating iCoh increase at stimulus onset, the median was calculated not only for the frequency range, but also across time in the first 100 ms after the sequence onset.

### 4.7 LFP onset power analyses

To test to what extent coherent gamma oscillations in the FAF-AC network could be attributable to a broadband evoked response in the LFP, we explored the statistical dependence of gamma power on a trial by trial basis. Spectral properties were obtained using three different temporal windows (**Fig. 6**): *pre* (−110 ms – −20 ms relative to stimulus onset), *onset* (0 – 90 ms relative to stimulus onset), *late* (90 – 180 ms relative to stimulus onset), and *full* (0 – 180 ms relative to stimulus onset). All analyses were performed using the 5.28 Hz syllabic train, thereby guaranteeing that responses to only one syllable (i.e. the first syllable presented) were considered in the time windows of choice. All spectra were calculated using the Chronux toolbox, with 2 tapers and a TW product of 2, on a trial-by-trial basis. Power spectra were compared for every penetration, per trial, statistically probing changes in the power of low-frequency (0-15 Hz) and gamma bands (25-45, 45-80, and 25-80 Hz) in the *pre* vs. *onset* windows (**Fig. S2A**; Wilcoxon signed rank tests, significance when p < 0.01), as well as during the *pre* vs. *late* periods (**Fig. 6C**). A percentage of significant difference (ratio across 50 penetrations) is depicted in the abovementioned figures. The time-frequency analyses shown in **Fig. 6D** was obtained by evaluating significance differences, per penetration, at given time windows (90 ms length) which were slid (10 ms steps) over times surrounding stimulus onset. Each time window spectra were compared, on a trial-by-trial basis given a penetration, with a window located before stimulus onset (center at ca. −90 ms relative to onset; Wilcoxon signed rank tests, significance when p < 0.01).

As per Medvedev and Kanwal (2008), the power spectra from each penetration were z-normalized across trials, and the power in a given band was calculated by integrating (*trapz* function, Matlab) over the z-normalized spectrum (per trial). Care was taken that the number of frequency samples were comparable when integrating at different bands; the gamma band was therefore divided into 25-45 and 45-60 Hz sub-bands. We then determined whether the distribution of power in gamma and low-frequencies were different, per penetration, by means of a 2-sample Kolmogorov Smirnov test (alpha at 0.01). Because differences or lack of differences in the power distributions do not necessarily imply the existence (or lack) of trial-by-trial correlation between the power of low and gamma frequency bands, we tested whether these powers were correlated for every penetration, on a trial by trial basis. In this context, correlations were significant given a p < 0.01.

### 4.8 Information theoretic analyses

Information in the neuronal response regarding the acoustic stimuli was quantified by means of Shannon’s mutual information (MI; (Shannon, 2001)). The MI between a stimulus set *S* and a response set *R* is mathematically expressed as follows:

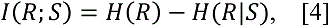

where *H(R)* is the response entropy (i.e. the overall variability of the response set), which is expressed as:

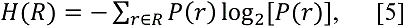

while

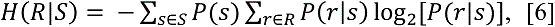

is referred to as the “noise entropy”, representing the irreproducibility of the response given a stimulus. The probabilities *P(r)*, *P(s)* and *P(r|s)* indicate the probability of observing response *r* taken from the set *R*, the probability of observing stimulus *s* from the set *S*, and the probability of observing response *r* given stimulus *s*, respectively. If the logarithm in **Eqs. 5** and **6** is of base 2, the MI has units of bits. Each bit of information means that an external observer is able to reduce, by observing the response, the uncertainty about the stimulus by a factor of 2 on a single trial basis. These quantities were estimated by means of the Information Breakdown Toolbox (ibTB; (Magri et al., 2009)).

#### 4.8.1 Stimuli for MI computations

With aims of quantifying the amount of information provided by FAF and AC spiking regarding a specific acoustic stream, we calculated each unit’s ability to discriminate consecutive chunks of the stimulus from each other (de Ruyter van Steveninck et al., 1997; Kayser et al., 2009; Kayser et al., 2010; Garcia-Rosales et al., 2018a). A particular sequence *S* (be it, for example, the natural the distress call) was subdivided into non-overlapping, consecutive segments *s_k_* (k = 1, 2, 3, …, M), all of length T = 4 ms. We chose this segment length so that results from this paper would be comparable with previous data from the AC of *C. perspicillata* (Garcia-Rosales et al., 2018a). Using lengths in the range of 2-12 ms did not alter the results qualitatively. Each segment *s* was treated as an independent substimulus from the set *S*. Note that, in this framework, all substimuli are equiprobable.

#### 4.8.2 Rate and joint neuronal codes

The manner in which *P(r)* is quantified depends directly on the assumptions made to characterize the neuronal response (i.e. the neural code considered). Here we used a rate code (I_rate_), which determines how well a unit discriminates between each substimulus *s*, based on its spiking rate. The response set represented whether a spike occurred or not, and can be characterized as follows: *R = {0, 1}*, where 1 and 0 represent the occurrence or absence of a spike, respectively. *P(r)* was then the probability that a unit fired or not a spike across all trials, whereas *P(r|s)* was the probability of firing to a certain substimulus. The time window was sufficiently short to assume that, in general, a single spike would occur within each time segment, and therefore binarized responses were used for MI calculations.

The information provided by joint responses from the FAF and the AC (I_joint_) was calculated by taking into account which unit elicited a spike in a merged response (see **Fig. 6B**; also referred to as “line code” in the literature (Panzeri et al., 2007; Kayser et al., 2009)). That is, the response set was defined as *R = { (0, 0); (0, 1); (1, 0); (1, 1) }*, where each member of the set represents whether and which unit fired a spike (e.g. *(0, 1)* could indicate that the FAF unit did not fire, whereas the AC one did; *(1, 0)* would represent the opposite).

#### 4.8.3 Quantifying information from limited samples

The probabilities in **Eqs. 5** and **6** are estimated empirically from the data, based on the representation of neuronal responses described above (i.e. the neural codes). These empirically estimated probabilities (such as *P(r)*, or *P(r|s)*) are biased because it is impossible in practice to sample all possible values of *R* a sufficiently large number of times (ideally, infinite). A number of methods have been developed to deal with the sampling bias (Panzeri et al., 2007). In this study, we used the Quadratic Extrapolation (QE) procedure (Strong et al., 1998), implemented in the ibTB. In addition to the QE, we subtracted possible remaining biases by means of a bootstrap procedure (Montemurro et al., 2008; Garcia-Rosales et al., 2018a), using 250 repetitions. For paired responses, we also used the shuffling procedure (Panzeri et al., 2007) implemented in the iBTB together with the bootstrap method. To corroborate that the information estimates presented in the results were not affected by the limited sampling bias, we conducted numerical simulations in order to measure the dependence of the bias on the number of trials. The results of these simulations are shown in Fig. S8, and indicate that the number of trials used in this study (50) was sufficient to robustly estimate the information quantities presented.

### 4.9 Statistical analyses

All statistical analyses were conducted in Matlab (version 8.6.0.267246 (R2015b)), with custom-written scripts using the Statistics and Machine Learning Toolbox. Tests for comparisons between the distributions of the quantities described above were always indicated in the main text. When multiple comparisons were done, we performed False-Discovery Rate (FDR) corrections (e.g., comparing across multiple channel pairs in **Fig. 3E**) with the Benjamini and Hochberg procedure (Benjamini and Hochberg, 1995). The significance threshold was set at an alpha of 0.05. If the p values reported were uncorrected, it is stated so in the text. Effect sizes were calculated with the *r* metric, which is defined as follows (Fritz et al., 2012):

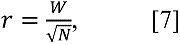

where *r* is the effect size, *W* is the test statistic of the Wilcoxon signed rank test used in this context, and N is the sample size of the quantities being compared (N = 50). Values of *r* ≤ 0.3 were considered small effects, while 0.3 ≤ *r* ≤ 0.5 were considered as medium effects, and large effects were considered when r > 0.5 (Fritz et al., 2012).

## 6 Author Contributions

**F.G.R** and **J.C.H** designed the study. **F.G.R** collected the data. **F.G.R** analyzed the data and wrote the manuscript. **F.G.R**, **L.L.J**, **E.G.P**, **Y.C.C**, and **J.C.H** discussed the results and reviewed the manuscript.

## 7 Acknowledgments

We are thankful to Manfred Kössl for comments on an earlier version of this manuscript.

## 1 Data Availability Statement

The data that support the findings of this study are available from the corresponding authors upon reasonable request.

