## Supplemental data for "Fronto-temporal coupling dynamics during spontaneous activity and auditory processing"

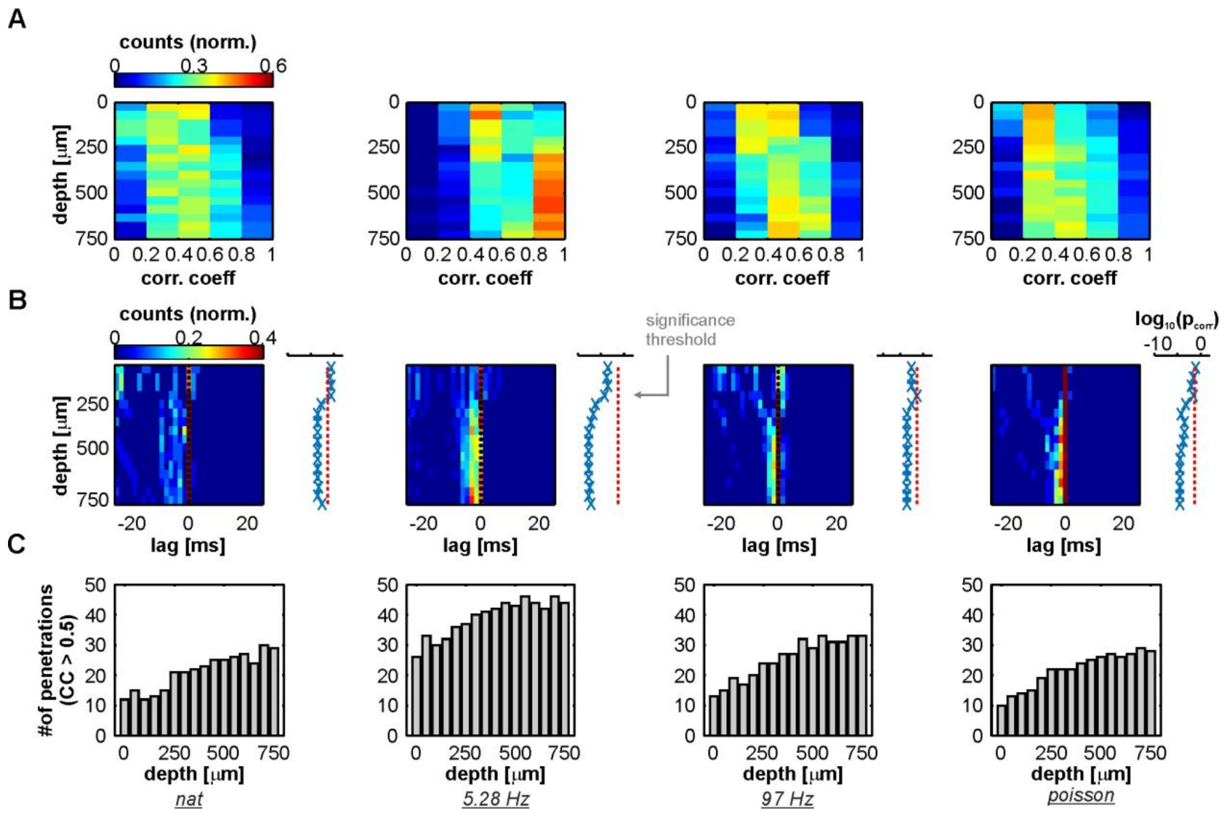


**Fig. S1. Well-correlated LFP stimulus-related activity in the FAF precedes that of the AC.** (Related to **Fig. 1**). (**A**) Distribution of correlation coefficients between simultaneously recorded LFPs in FAF and AC at various depths (N = 50 penetrations; stimulus left to right: natural call, 5.28 Hz train, 97 Hz train, and Poisson train). (**B**) Heatmaps: distribution across stimuli of cross-correlation lags between LFPs from FAF and AC, using only LFP pairs whose correlation coefficient was > 0.5. FDR corrected p values resulting from comparing the distributions, per AC depth, with a median of 0 (Wilcoxon signed rank test, significance for $p_{corr} < 0.05$). (**C**) Number of observations, per stimulus and AC depth, for which the correlation coefficient between LFPs in both structures were > 0.5 (equivalent to the sample size for statistical comparisons in panel **B**). In the figure, each column corresponds to the analysis for a different stimulus (indicated at the bottom). Note that observations regarding LFPs in AC "lagging" those in FAF are robust, even when considered well-correlated traces.

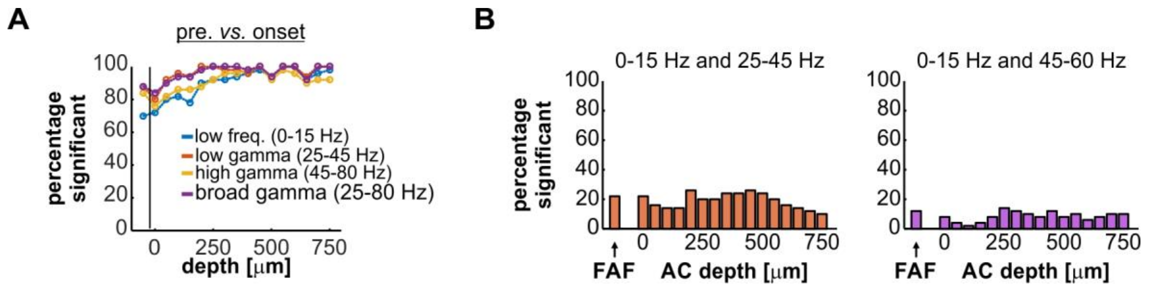


**Fig. S2.** (Related to **Figs. 6 and 7**). (**A**) Percentage of penetrations (from a total of 50) for which there were significant differences, for various frequency bands (indicated in the figure), between the power during the *onset* period (segment of 90 ms directly after stimulus onset) and the *pre* period (segment of 90 ms before stimulus onset) of the 5.28 Hz stimulus train (see **Fig. 6** in the main text). (**B**) Percentage of penetrations for which there was a significant correlation between the relative power of gamma (at bands of 25-45 and 45-60 Hz) and low-frequencies at the *full* period (see **Fig. 6**). These distributions complement the data shown in **Fig. 7**.

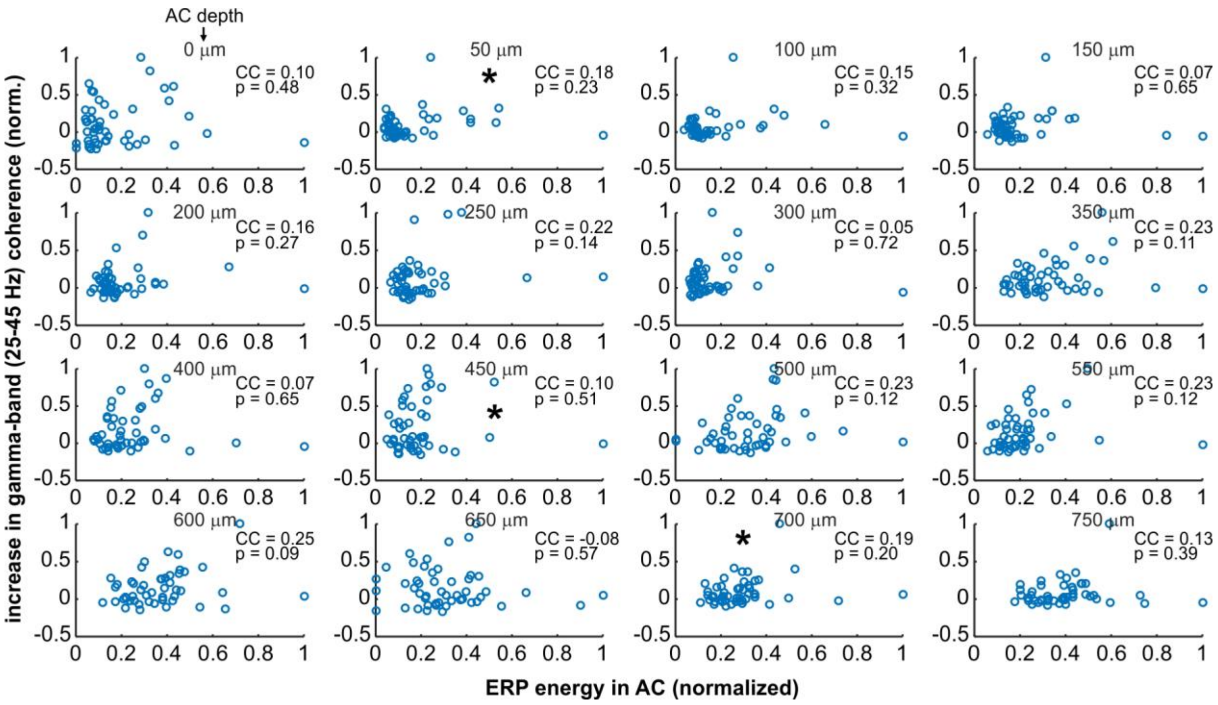


**Fig. S3. Correlation between event-related potential energy in the AC (ERP energy) and the gamma-band coherence increase in the FAF-AC circuit.** (Related to **Figs. 5, 6, 7**). Each shows a scatter plot (at different AC depths, indicated in the figure) of the energy in the event-related potential and the increase in gamma-band coherence, for each penetration tested (n = 49). ERP energy was obtained as the area under the average Hilbert transform (absolute value) across trials, per penetration, in a time window of 0-150 ms after stimulus onset (same period used to estimate coherence increase in the main text). Data of gamma-band coherence increase is the same shown in **Fig. 5**. Note the lack of significant correlation, across AC channels, between ERP energy and gamma-band coherence increase. Panels marked with an asterisk (*****) correspond to data shown in **Fig. 7I**.

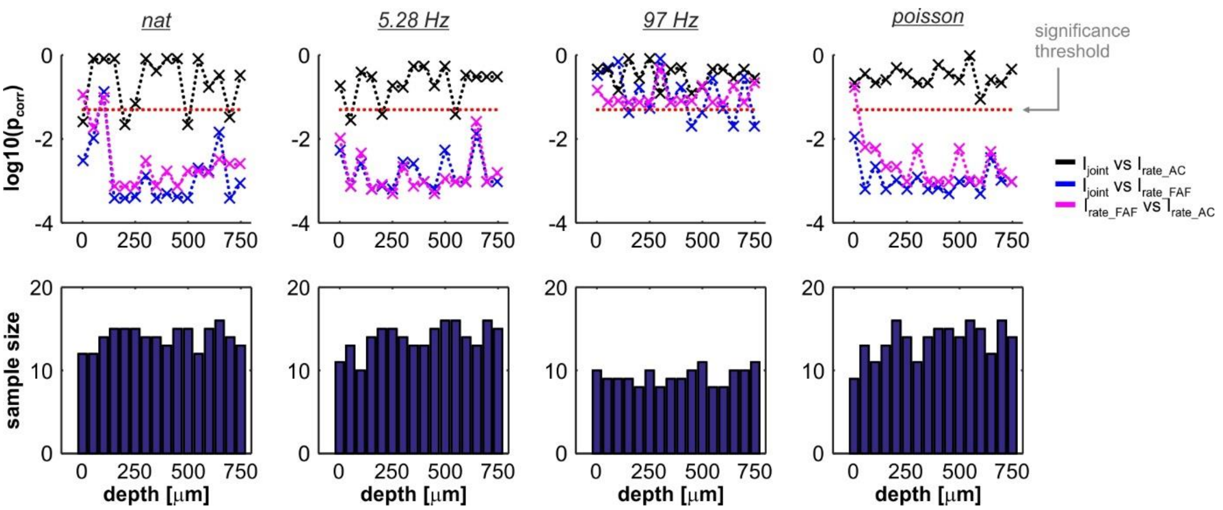


**Fig. S4. Comparison between paired $I_{joint}$, $I_{rate\_AC}$ and $I_{rate\_FAF}$.** (Related to **Fig. 8**). *Top*: Corrected p values resulting from comparing $I_{joint}$ and $I_{rate\_AC}$ and $I_{rate\_FAF}$ calculated from paired responses (FDR corrected Wilcoxon signed rank tests, significance when $p_{corr} < 0.05$, indicated as a red trace in the panels; $I_{joint}$ vs. $I_{rate\_FAF}$, blue; $I_{joint}$ vs. $I_{rate\_AC}$, black; $I_{rate\_AC}$ vs. $I_{rate\_FAF}$, magenta). Comparisons were performed across stimuli and depths in the AC. Note that $I_{rate\_FAF}$ was typically lower than $I_{rateAC}$, and $I_{joint}$. There were very few instances of significant differences between $I_{rate\_AC}$ and $I_{joint}$. *Bottom*: Sample size of the populations used to calculate the p values shown in the top panels. Sample sizes below 10 in the case of the 97 Hz may have affected statistical outcomes, particularly after correcting for multiple comparisons. These sample sizes are equivalent to the data shown in **Fig. 6C**.

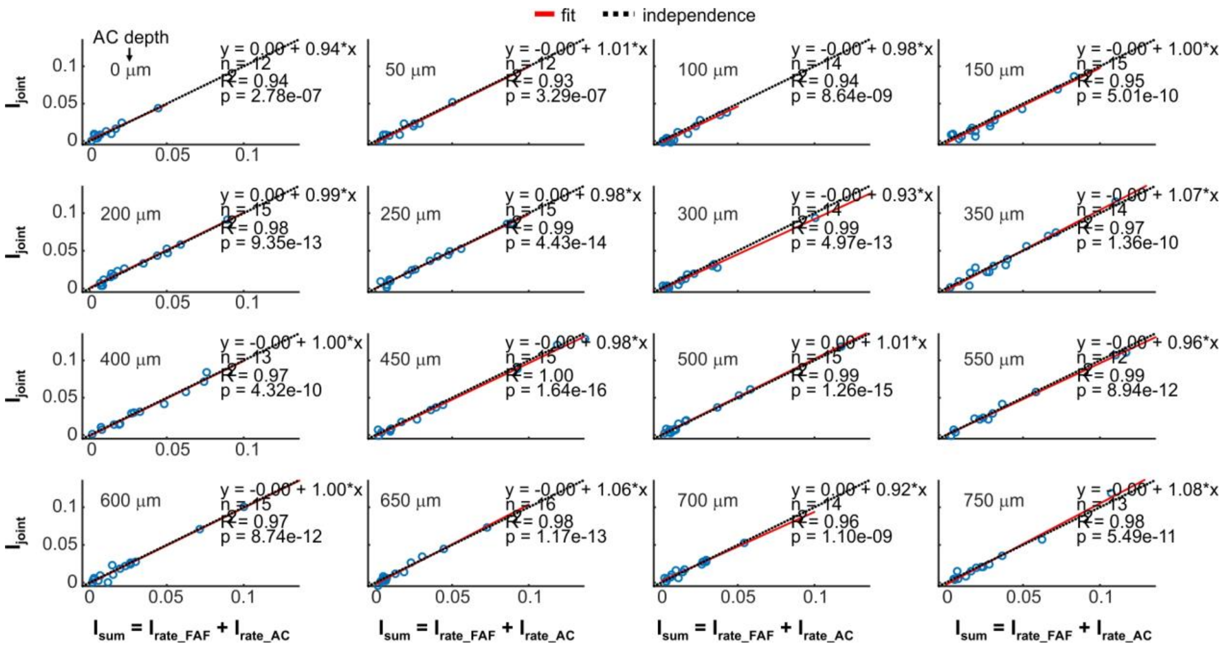


**Fig. S5. Relationship between $I_{sum}$ and $I_{joint}$ during the processing of the natural distress sequence.** (Related to **Fig. 8**). Each graph shows the relationship between $I_{sum} = I_{rate\_FAF} + I_{rate\_AC}$ (that is, the sum of the $I_{rate}$ of each unit conforming the FAF-AC pair), during the processing of the natural sequence. A red line indicates the linear fit, whereas a black dashed line marks the regime of independence in the plots. Graphs were obtained from each channel in the AC (depth indicated by the upper left legend), together with the FAF. On the upper right region of the graphs, information is given regarding the curve of the linear fit (y = ax + b), the number of points used (n), the coefficient of determination ($R^2$), and the p value of the fit (p).

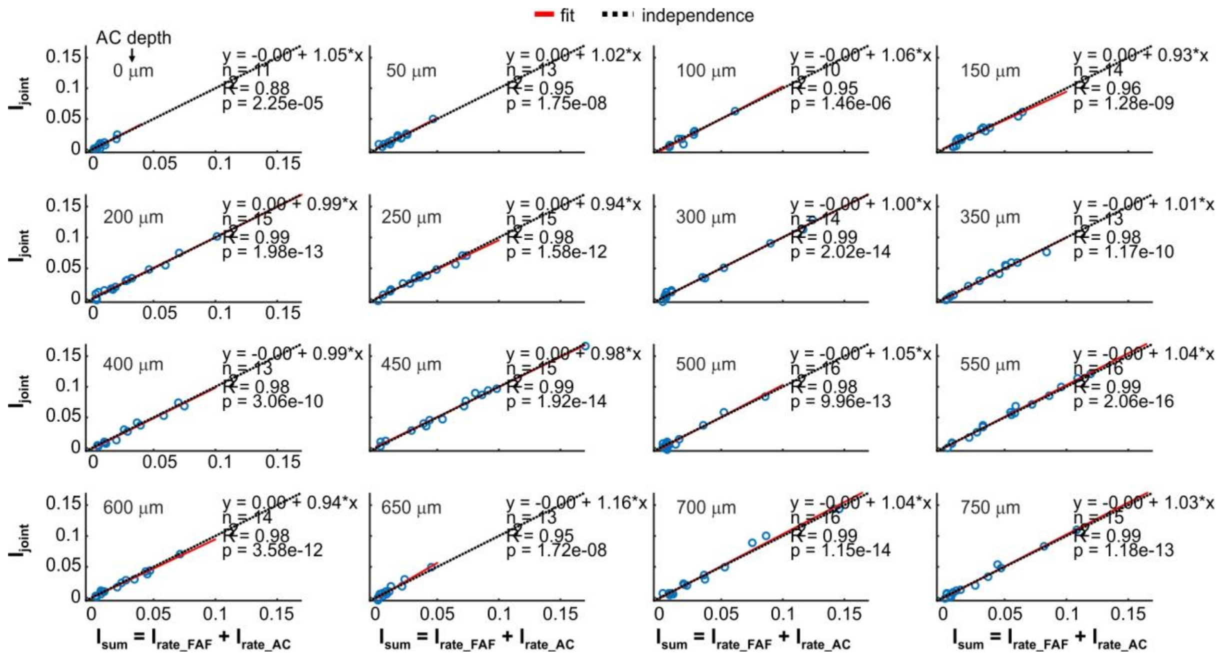

**Fig. S6. Relationship between $I_{sum}$ and $I_{joint}$ during the processing of the 5.28 Hz syllabic train.** (Related to **Fig. 8**). This figure follows the conventions from **Fig. S5**, but data was acquired during the processing of the 5.28 Hz syllabic train.

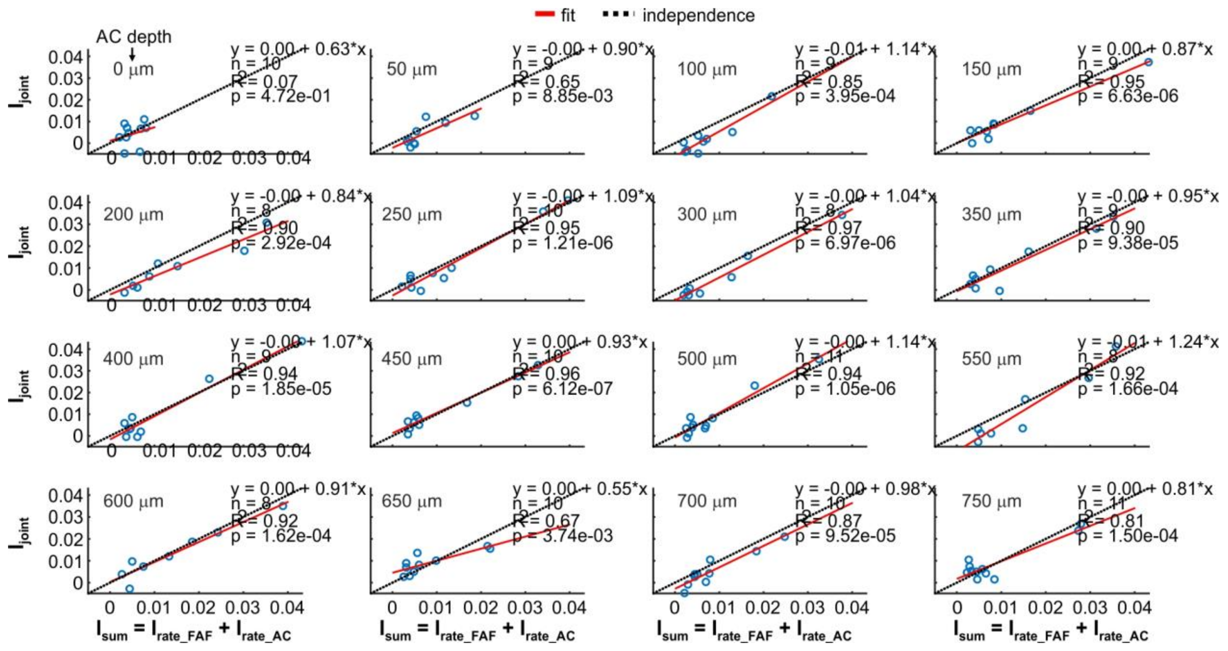

**Fig. S7. Relationship between $I_{sum}$ and $I_{joint}$ during the processing of the 97 Hz syllabic train.** (Related to **Fig. 8**). This figure follows the conventions from **Fig. S5**, but data was acquired during the processing of the 97 Hz syllabic train.

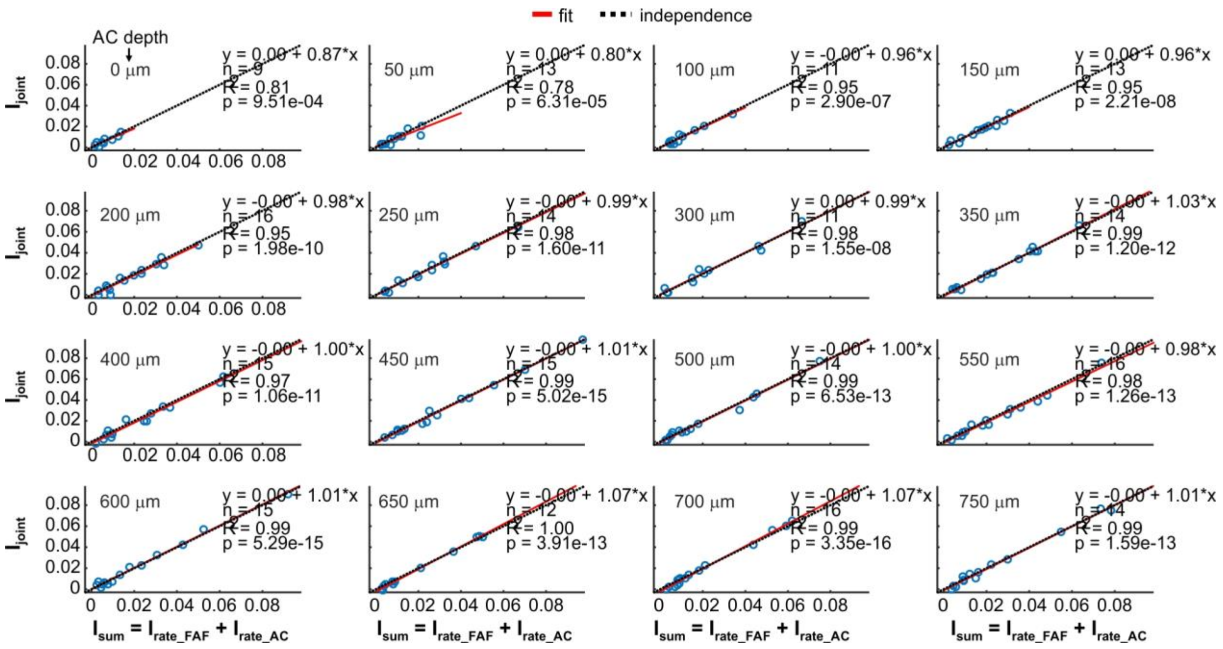


**Fig. S8. Relationship between $I_{sum}$ and $I_{joint}$ during the processing of the Poisson syllabic train.** (Related to **Fig. 8**). This figure follows the conventions from **Fig. S5**, but data was acquired during the processing of the Poisson syllabic train.

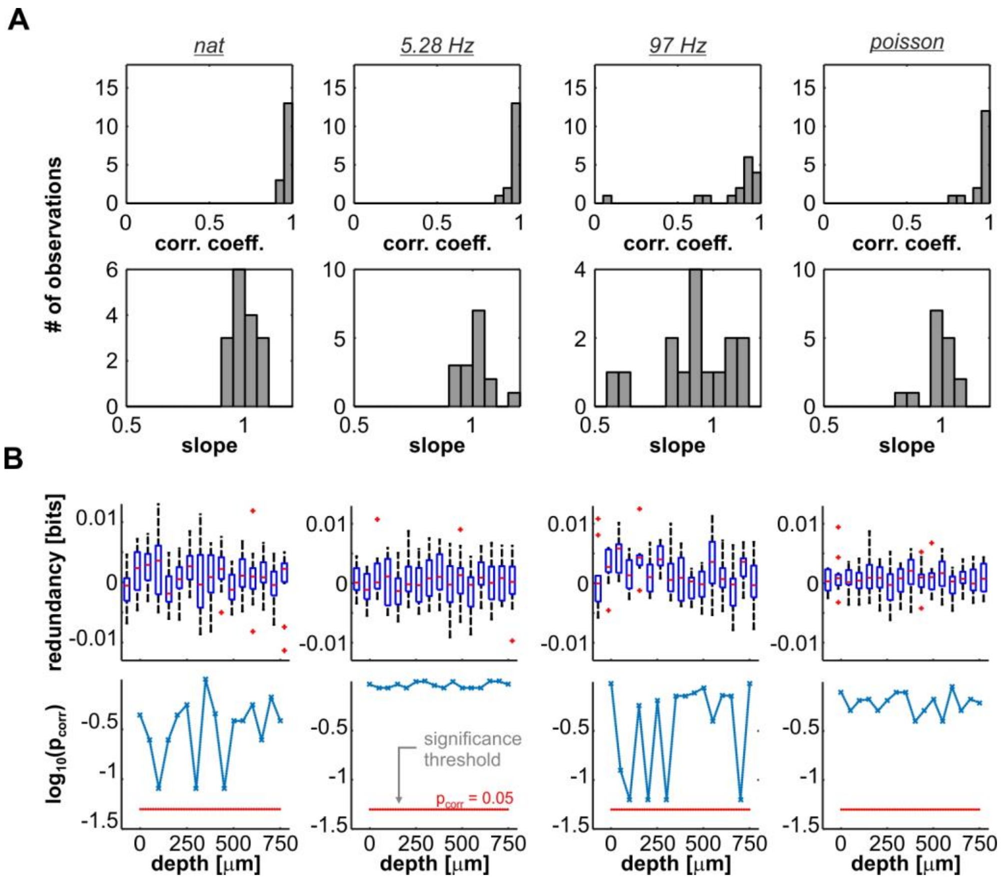


**Fig. S9 Summary of linear dependence between $I_{sum}$ and $I_{joint}$.** (Related to **Fig. 8**). (**A**) Top: coefficients of determination ($R^2$) as calculated from data in **Figs. S5-8** for all channels in response to the acoustic sequences. High $R^2$ values were typical. Bottom: slope of the linear fits depicted in **Figs. S5-8**, across stimuli, for all channels. Note that the majority of the slopes are close to 1. (**B**) Top: Redundancy as quantified from the difference between $I_{sum}$ and $I_{joint}$. Positive values indicate response redundancy in FAF-AC spiking; negative values indicate synergistic interactions; while values of 0 suggest independence. The distributions across channels were not statistically different from zero (bottom; FDR-corrected Wilcoxon signed rank tests, $p_{corr}$ values indicated in the panels, per stimulus, in a logarithmic scale), but the former does not necessarily imply a general trend towards independence in our dataset (see **Fig. 8E**).

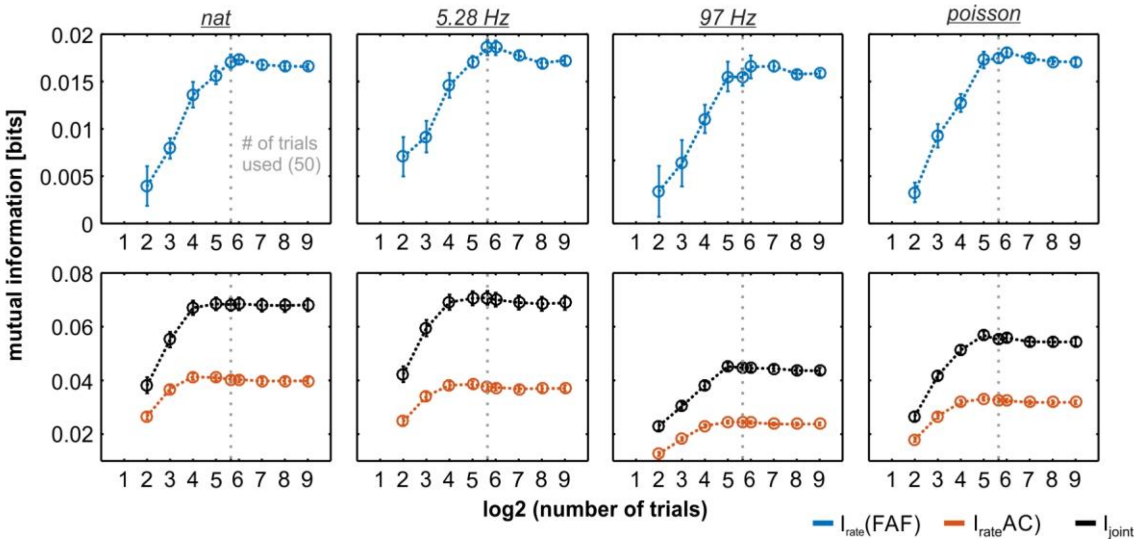


**Fig. S10 Performance of bias correction for information theoretic calculations**. (Related to **Fig. 8**). In order to evaluate the performance of the bias corrections in our information theoretic calculations, we simulated spike trains whose first-order statistics were close the those of the real data. The number of trials available from the synthetic dataset was systematically varied to explore the effect of trial number on the estimation of mutual information provided by each code considered in the study. The upper row of the figure shows, per stimulus, the $I_{rate}$ estimations from a dataset mimicking the FAF spiking, using different number of trials (4, 8, 16, …, 512; note that x-axis is in a logarithmic scale). The bottom row depicts similar comparisons of $I_{rate}$ from the AC (orange) and $I_{joint}$ (black), obtained with the synthetic dataset. Data from all channels were pooled when the AC was considered (for $I_{rate}$ and $I_{joint}$). The bias was negative for a number of trials lower than 32, and became negligible already for the number of trials used in the experiments (50; indicated with a vertical grey dashed line). In the figure, neuronal codes are calculated using a time window of 4 ms, and with binarized spike trains (same parameters used for the experimental data shown in the **Fig. 8**).
